# Plasma membrane nanodeformations promote actin polymerisation through CIP4/CDC42 recruitment and regulate type II IFN signaling

**DOI:** 10.1101/2022.08.16.504113

**Authors:** Ledoux Benjamin, Zanin Natacha, Yang Jinsung, Coster Charlotte, Dupont-Gillain Christine, Alsteens David, Morsomme Pierre, Renard Henri-François

## Abstract

In their environment, cells have to cope with mechanical stresses constantly. Among those, nanoscale deformations of plasma membrane induced by substrate nanotopography are now largely accepted as a biophysical stimulus influencing cell behaviour and function. However, the mechanotransduction cascades involved and their precise molecular effects on cellular physiology are still poorly understood. Here, using homemade fluorescent nanostructured cell culture surfaces, we explored the role of <u>B</u>in/<u>A</u>mphiphysin/<u>R</u>vs (BAR) domain proteins as mechanosensors of plasma membrane geometry. Our data reveal that distinct subsets of BAR proteins bind to plasma membrane deformations in a membrane curvature radius-dependent manner. Furthermore, we show that membrane curvature promotes the formation of dynamic actin structures mediated by the Rho GTPase CDC42, the F-BAR protein CIP4 and the presence of PI(4,5)P_2_, independently of clathrin. In addition, these actin-enriched nanodomains can serve as platforms to regulate receptor signaling as they appear to contain Interfero<u>n γ</u> receptor (IFNγ-R) and to lead to the partial inhibition of IFNγ-induced Janus-<u>a</u>ctivated tyrosine <u>k</u>inase/<u>s</u>ignal transducer and <u>a</u>ctivator of transcription (JAK/STAT) signaling.

## Introduction

In their environment, cells constantly experience a variety of external mechanical stimuli and stresses inducing membrane curvature and cell deformation, which they must respond to accordingly. Among them, cell deformation due to the presence of topographical features on the scale of hundreds of nanometers has been shown to alter key cellular functions such as adhesion [1,2], differentiation [3– 5], proliferation [6,7] and migration [8]. In parallel, plasma membrane curvature has been shown to influence several molecular processes, including focal adhesion formation [1,8,9], clathrin-mediated endocytosis [10,11] and actin polymerisation [12,13]. However, numerous gaps remain in the understanding of these molecular mechanisms and their connection with changes in cellular behaviour. In particular, it is still not known how cells are able to sense and then distinguish between various types of deformations in terms of size, geometry and dynamics leading them to finely tailor their responses.

Thanks to their properties, it is not surprising to find <u>B</u>in/<u>A</u>mphiphysin/<u>R</u>vs (BAR) domain proteins as main actors of these dynamic molecular machineries. Indeed, multiple recent studies have shown that BAR domain proteins play pivotal roles in membrane geometry recognition and processing [10,12,14– 17]. Importantly, BAR domain proteins form dimers featuring a crescent shape with a unique intrinsic curvature radius, allowing them to induce and/or bind membrane deformations of different geometries. They are classified into several subfamilies mostly based on their structures and associated domains. Generally, N-BARs (containing an N-terminal amphipathic helix) harbour a highly curved structure while Fes/CIP4 homology BARs (F-BAR) are shallower [18–20]. Then, after binding membranes through their BAR domain, they can promote local molecular responses as they often feature additional functional domains such as <u>S</u>rc <u>h</u>omology <u>3</u> (SH3) or Rho <u>G</u>TPase <u>a</u>ctivating <u>p</u>rotein (GAP) domains. In particular, F-BAR domain proteins from the <u>T</u>ransducer <u>o</u>f <u>C</u>DC42-dependent <u>a</u>ctin assembly (TOCA) subfamily (CIP4, FBP17 and FNBP1L) have been linked to the remodeling of the actin cytoskeleton through activation of N-WASP and Arp2/3 in response to topography-induced membrane curvature [12]. Yet, the regulation and the purpose of this actin reorganisation are still largely unknown. Interestingly, CIP4 and FBP17 together with the Rho GTPase CDC42 have also been involved in the priming of <u>F</u>ast <u>E</u>ndophilin-<u>M</u>ediated <u>E</u>ndocytosis (FEME) carriers, another membrane curvature-driven process, independent of clathrin [21].

In this study, we developed a simple colloidal lithography technique to synthesise fluorescent nanostructured substrates which were used to induce spherical and isotropic nanoscale plasma membrane deformations ranging from 100 nm to 500 nm in living cells. Using this tool in combination with high-resolution confocal microscopy, we showed that a distinct subset of BAR domain proteins was recruited to each plasma membrane deformation size. Interestingly, the affinity of BAR proteins for a particular curvature radius does not correlate with their usual subfamily classification. Our data further demonstrate that membrane curvature promotes the formation of dynamic actin structures mediated by the Rho GTPase CDC42 (and not Rac1), the F-BAR protein CIP4 and the presence of PI(4,5)P_2_, independently of clathrin. Furthermore, using correlative FluidFM/fast-scanning confocal microscopy, we demonstrated that these actin structures can form both at basal and apical plasma membranes, in a very dynamic manner. In addition, these membrane nanodomains are reminiscent of early-stage FEME-like endocytic pits, opening up the exciting possibility of the existence of frustrated clathrin-independent endocytosis, as has been shown with the frustrated clathrin-mediated endocytosis previously described [22–24]. Furthermore, these actin-enriched nanodomains harbour Interfero<u>n γ</u> receptors (IFNγ-R) with an impaired signaling. Altogether, our data demonstrate that plasma membrane deformation induces the formation of CDC42/CIP4/PI(4,5)P_2_-dependent actin structures which can serve as platforms to regulate receptor signaling.

## Results

### A versatile approach to generate nanoscale plasma membrane deformations

Multiple nanofabrication techniques exist to create a variety of nanopatterns used for the study of cellular responses to membrane deformations. While some are being limited by their complexity and/or low throughput (*e*.*g*. electron beam lithography), others are relatively simple to work with. Here, we took advantage of both the rapidity and simplicity of colloidal lithography to produce substrates featuring fluorescent spherical nanostructures [25]. Briefly, negatively charged spherical silica colloids were coated with a positively charged polymer, polyallylamine (PAH) (**Figure 1a**). The added amine groups were then functionalized using succinimidyl ester conjugated fluorescent dyes (ATTO 390, Alexa Fluor 488, Alexa Fluor 555 or ATTO 647N), leading to the generation of fluorescent colloids. Next, these colloids were adsorbed onto a glass coverslip to create the nanostructures (**Figure 1a**). Finally, the substrate was coated with collagen to make the surface chemistry uniform with stable nanostructures and appropriate cell adhesion properties (**Figure 1a**). This approach allowed the quick and cost-effective patterning of relatively large surfaces for cell culture (12- and 30-mm round coverslips) with fluorescent nanostructures of 500 nm, 300 nm and 100 nm in diameter at various densities (**Figure 1b-c**). Despite their small size and random distribution, the nanostructures are easily distinguished on the substrate thanks to their fluorescence. In addition, in contrast to commercial fluorescent polystyrene beads, our nanostructures have the advantage of displaying both the same surface chemistry and stiffness as the underlying glass substrate, avoiding potential effects due to substrate heterogeneity.

**Figure 1:**
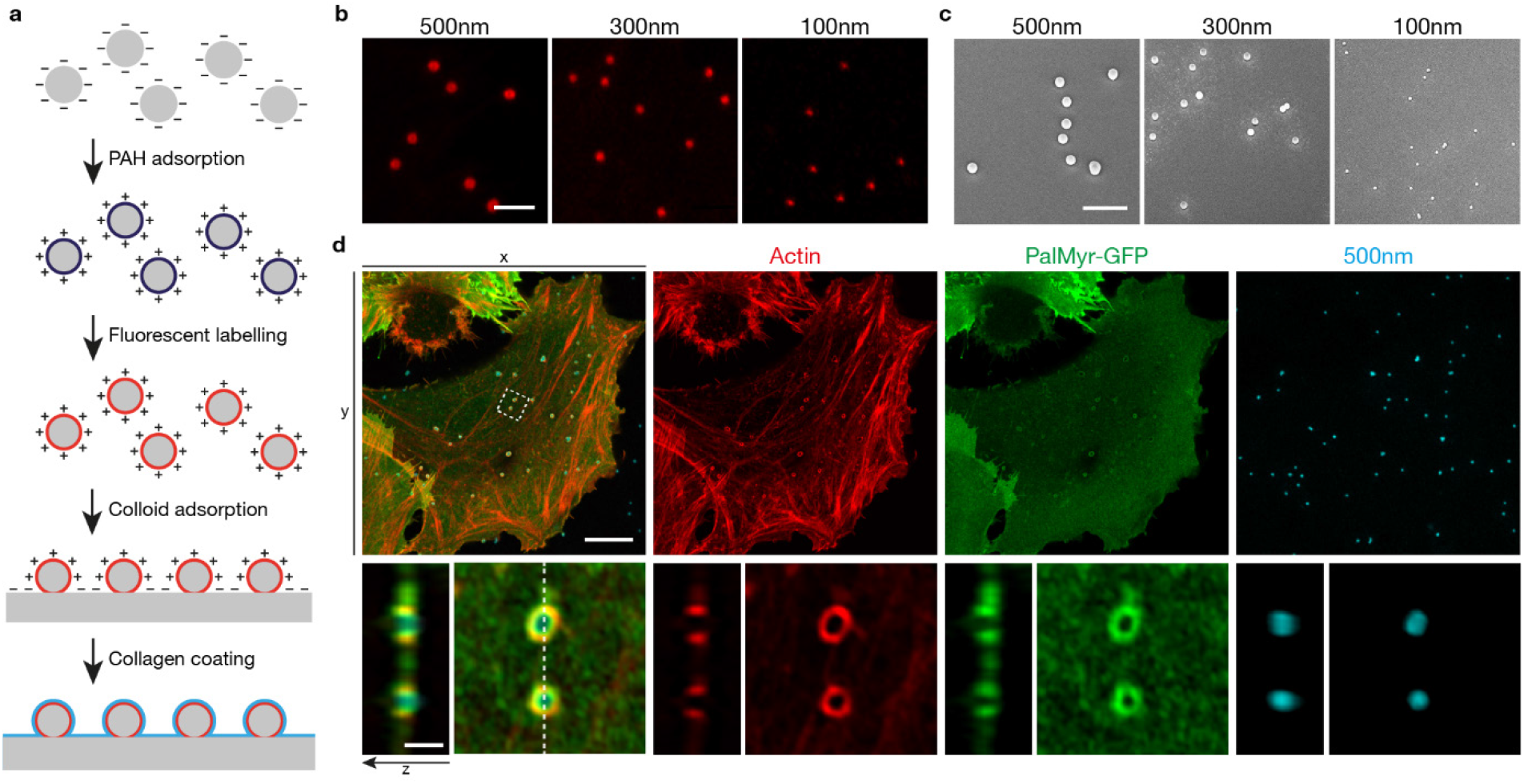
Nanostructures deform cell plasma membrane and induce local actin polymerisation. **a**, Synthesis of nanostructured substrates. Silica colloids were coated with polyallylamine (PAH), labeled with fluorescent dyes and adsorbed onto coverslips. Coverslips were then coated with collagen before seeding cells. **b-c**, ATTO 390 labeled nanostructures of 500 nm, 300 nm and 100 nm visualized by Airyscan confocal microscopy (**b**) or SEM (**c**). **d**, HeLa cells expressing PalMyr-GFP (green) seeded on 500 nm nanostructures (cyan) and stained for actin (phalloidin, red). Representative image from > 3 independent experiments (**b,d**) or one single experiment (**c**). Region marked by a dashed square, expanded below (panel **d**, bottom) and also displayed as a transversal view along Z-axis. Representative images from > 3 independent experiments (**b,d**) or a single experiment (**c**). Scale bars, 10 µm (**d**, top), 2 µm (**b-c**) and 1 µm (**d**, bottom).

When cells were seeded on nanostructured substrates, their plasma membrane deformed and wrapped around the nanostructures. This resulted in a local deformation of the plasma membrane with a curvature radius determined by the diameter of the nanostructure used. This wrapping was visualized using high-resolution Airyscan confocal microscopy by expressing a palmitoylated and myristoylated (PalMyr) GFP as a plasma membrane marker (**Figure 1d**, green channel). Additionally, the actin cytoskeleton reacted to this curvature and underwent reorganisation, forming actin rings (**Figure 1d**, red channel). Both membrane wrapping and actin ring structures were observable on nanostructures of 500 nm and 300 nm (**Figure 1d, Supplementary Figure 1a**). However, due to the resolution limit of Airyscan microscopy, we were only able to observe local actin polymerisation in the form of dots on 100 nm nanostructures (**Supplementary Figure 1b**). Nevertheless, using more highly resolved STED (<u>St</u>imulated-<u>E</u>mission-<u>D</u>epletion) nanoscopy, we confirmed that those dots were indeed unresolved actin rings (**Supplementary Figure 6**). Although the nanostructures are not covalently bound to the surface, they were mostly stable and neither displaced nor internalized by cells for a minimum of 2 hours after cell seeding (**Movie 1**). Together, these data underline the relevance of our nanostructured substrate to study biomolecular mechanisms triggered by the bending of the plasma membrane.

#### BAR domain proteins bind to plasma membrane deformations in a membrane curvature radius-dependent manner

To investigate how cells can detect and respond to nanoscale curvature of their plasma membrane, we performed a quantitative screening, focusing on BAR domain protein affinity towards membrane deformations of 500 nm, 300 nm and 100 nm (**Figure 2a-e**). Thanks to their curved shape and their ability to sense membrane curvature, BAR domain proteins are ideal candidates to connect plasma membrane geometry to downstream cellular responses. Of note, some of them have already been shown to be involved in such mechanisms [14,10,12,26]. To identify which BAR domain proteins could be potential candidates in mechano-transduction responses, HeLa cells transiently expressing fluorescently-tagged BAR domain proteins (69 constructs screened) were seeded on nanostructured substrates for 1 hour and fixed. Cells were then imaged using high-resolution Airyscan confocal microscopy, and the fluorescence around each individual nanostructure was quantified using a homemade (semi-)automated method (see **Supplementary Figure 2** and methods for details about the quantification procedure) (**Figure 2a-d**). As a negative control, we used PalMyr-GFP to take into account the non-specific gain in fluorescence due to the bending of the membrane perpendicularly to the focal plane. BAR proteins with an enrichment ratio significantly higher than PalMyr-GFP were considered as enriched, thus having a higher affinity for the corresponding deformation (**Figure 2a-d**).

**Figure 2:**
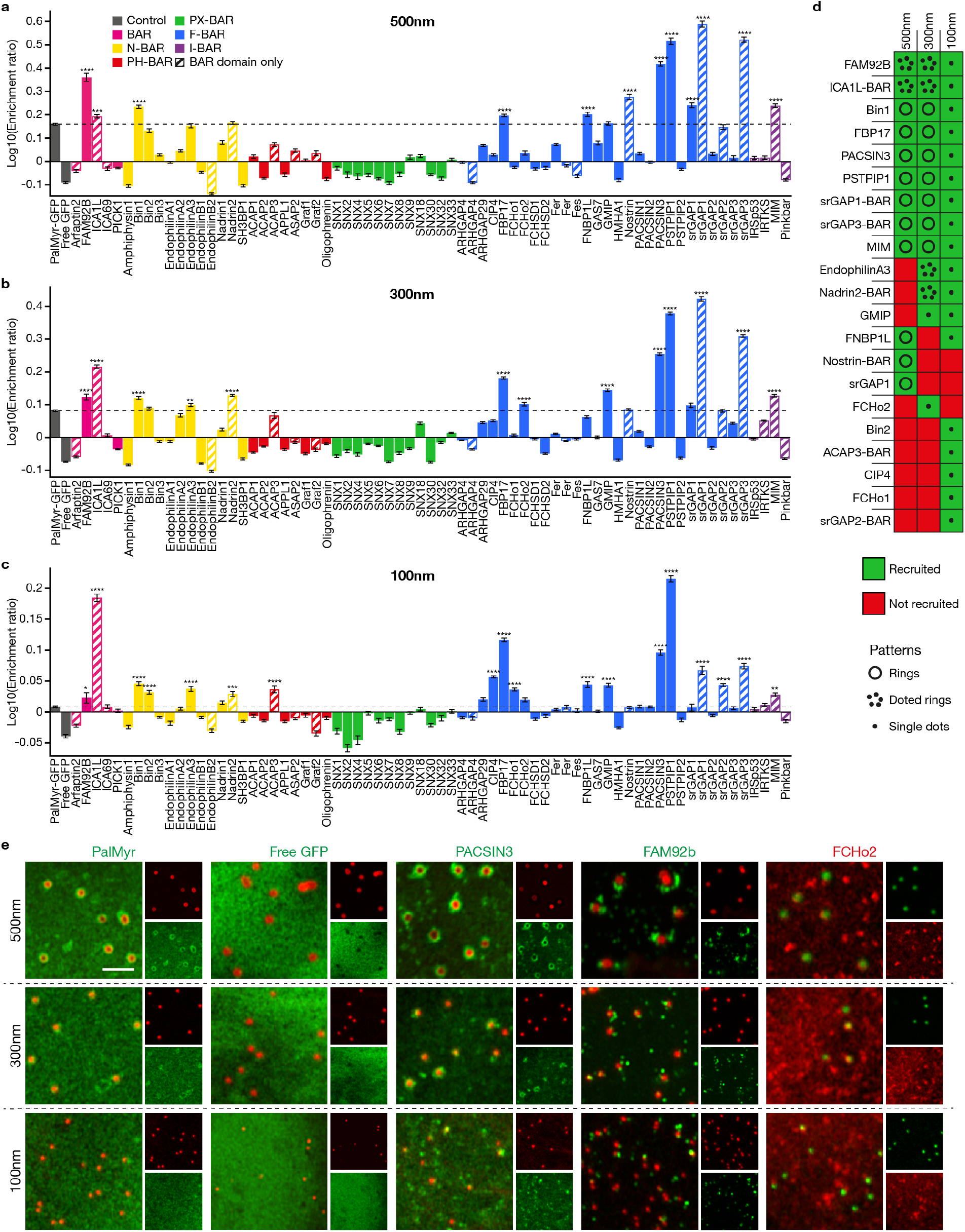
BAR domain proteins bind to plasma membrane deformations in a membrane curvature radius-dependent manner. **a-c**, Quantification of Airyscan images showing the recruitment of fluorescently-tagged BAR domain proteins to 500 nm (**a**), 300 nm (**b**) and 100 nm (**c**) plasma membrane deformations in HeLa cells (more details on the quantification procedure in **Supplementary Figure 2**). Data are mean ± s.e.m. from two independent experiments with a minimum of 158 deformations per construct (see **source data files** for exact *n* for each condition). Each protein was compared to the recruitment of PalMyr-GFP used as a control (dashed lines). **** P < 0.0001; *** P < 0.001; ** P < 0.01; * P < 0.05 (one-way ANOVA with Dunnett’s multiple comparison test). **d**, Summary table of recruited BAR domain proteins and their patterns. **e**, Representative images of controls (PalMyr-GFP and GFP) and selected BAR domain proteins depicting different patterns observed around membrane deformations. Representative images for each condition in **Supplementary Figure 3**. Scale bar, 2 µm (**e**).

Strikingly, we observed that different subsets of BAR proteins were recruited according to the size of the deformations (**Figure 2d**). Indeed, although some were recruited independently of the size of the deformation (*e*.*g*. PSTPIP1, PACSIN3), others displayed more specificity towards smaller/larger sizes (*e*.*g*. CIP4 towards 100 nm). However, some discrepancies in the local accumulation at the plasma membrane were observed between the fusion proteins expressing the full BAR protein sequence versus the BAR domain alone. Indeed, in the case of srGAP1-3, it resulted in the BAR domains being highly recruited to deformations, while recruitment was barely observable for the full-length proteins. This might be due to autoinhibition of the BAR domain by the SH3 domain, as it was previously shown in the case of srGAP2 [27]. In addition, according to our assumptions, we detected a local accumulation of BAR proteins which were previously studied for their curvature sensing ability such as Nadrin2, FCHo1, CIP4, FNBP1L and FBP17 [10,12,14]. Interestingly, we also observed the recruitment of an inverted-BAR (I-BAR) protein, MIM. It is most likely recruited to the base of the deformed membrane, where the curvature is negative.

Interestingly, we were able to distinguish three distinct patterns around deformations among the twenty-one hits at the resolution of Airyscan: uniform rings, single dots or cluster of dots surrounding individual deformations (**Figure 2d-e**). Surprisingly, the affinity of BAR domain proteins for a particular membrane curvature radius does not correlate with their intrinsic protein curvature radius, usually represented by their subfamily classification. Indeed, both F-BAR and N-BAR proteins were found around all curvature sizes tested. This observation may appear rather counterintuitive, as N-BAR domains are usually considered to favour much higher membrane curvature than F-BAR domains [18,19]. Altogether, these results indicate that cells are able to use distinct arsenals of BAR domain proteins to distinguish between different plasma membrane deformation geometries.

#### CDC42 drives the recruitment of CIP4 and polymerisation of the actin cytoskeleton around plasma membrane deformations

A closer look at the screening results (**Figure 2a-d**) revealed several hits which are already known to play a role in cytoskeleton reorganisation and Rho GTPase signaling (srGAPs, TOCAs, Nadrins, GMIP) [28]. In particular, TOCA proteins (CIP4, FBP17 and FNBP1L) are well-known effectors of CDC42 and seem to be involved in curvature-mediated actin polymerisation [12]. Yet, it remains unknown if CDC42 plays any role in this curvature-mediated remodeling of the actin cytoskeleton.

To answer this question, we first explored the effect of transient expression of wild-type CDC42 (CDC42-WT) as well as its constitutively active (CDC42-Q61L) or dominant negative (CDC42-T17N) mutants (**Figure 3a-d**). Interestingly, when wild-type CDC42 was overexpressed, we observed an increase in both actin polymerisation and CIP4 recruitment to 100 nm deformations (**Figure 3a-c**). Furthermore, upon expression of the active CDC42-Q61L mutant, the recruitment of actin and CIP4 was even more pronounced, with the appearance of actin structures around most deformations (**Figure 3a-c**). In addition, we observed that CDC42-WT and its active mutant are both recruited to the deformations (**Figure 3a,d**). On the other hand, neither CDC42-T17N mutant nor Rac1 had an effect on CIP4 and actin recruitment (**Figure 3a-c**). As expected, CDC42, CIP4 and actin were almost systematically found colocalising around the deformations (**Figure 3a-d**, white arrowheads). Similar results were observed with 300 nm and 500 nm deformations (**Supplementary Figures 4a-d and 5a-d**): the expression of CDC42-Q61L increased local actin polymerisation and CIP4 recruitment, while CDC42-T17N had the opposite effect. Nevertheless, the effect was more pronounced for 100 nm deformations, with the formation of big patches of CIP4 and actin near individual deformations. Using STED nanoscopy to circumvent the resolution limit of Airyscan, we confirmed that CIP4 and actin were indeed recruited to 100 nm deformations in control cells and that it was not a random colocalisation, (**Supplementary Figure 6**).

**Figure 3:**
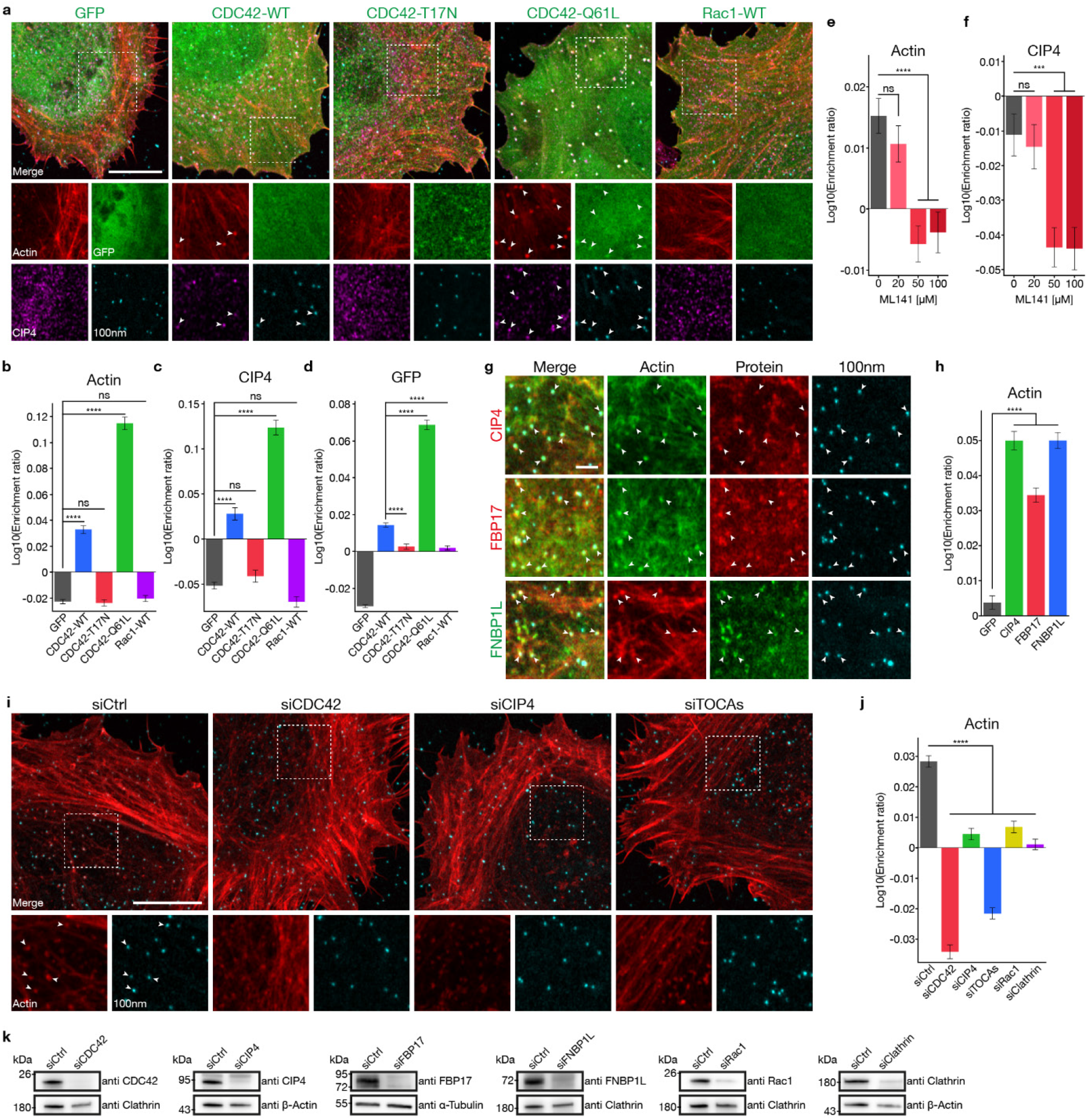
CDC42 and CIP4 control actin polymerisation around 100 nm plasma membrane deformations. HeLa cells grown on 100 nm nanostructures and transfected with fluorescent constructs (**a-d,g-h**), treated with a specific CDC42 inhibitor (ML141) (**e-f**) or with siRNAs (**i-j**), as indicated. Data shown are quantifications of actin (**b,e,h,j**), endogenous CIP4 (**c,f**) or GFP (**d**) fluorescence around 100 nm deformations and the corresponding representative Airyscan images (**a,g,i**), as indicated. **a-d**, Effect of transient expression of GFP, GFP-CDC42-WT, GFP-CDC42-T17N, GFP-CDC42-Q61L or GFP-Rac1-WT, on the enrichment of actin, CIP4 and GFP around deformations. Number of deformations: GFP, n = 8146; CDC42-WT, n = 3829; CDC42-T17N, n = 2993; CDC42-Q61L, n = 3058; Rac1-WT, n = 3942. Three independent experiments. **e-f**, Effect of dose-dependent inhibition of CDC42 by ML141 (0 µM, 20 µM, 50 µM or 100 µM) on the enrichment of actin and CIP4 around deformations. Number of deformations: 0 µM, n = 3007; 20 µM, n = 2541; 50 µM, n = 2788; 100 µM, n = 2252. Three independent experiments. **g-h**, Effect of transient expression of GFP or individual TOCA proteins (mCherry-CIP4, mCherry-FBP17 or GFP-FNBP1L) on the enrichment of actin around deformations. Number of deformations: GFP, n = 7285; CIP4, n = 5658; FBP17, n = 6220; FNBP1L, n = 5544. Four independent experiments. **i-j**, Effect of the depletion of CDC42, CIP4, TOCAs (CIP4, FBP17 and FNBP1L), Rac1 or Clathrin Heavy Chain with siRNAs on the enrichment of actin around deformations. Number of deformations: siCtrl, n = 6940; siCDC42, n = 3184; siCIP4, n = 6832; siTOCA, n = 7468; siRac1, n = 7552; siClathrin, n = 9056. Three independent experiments. **k**, Immunoblots against CDC42, CIP4, FBP17, FNBP1L, Rac1 and Clathrin Heavy Chain document siRNA knock-down efficiency. Data are mean ± s.e.m.; ns, not significant; **** P < 0.0001; *** P < 0.001 (one-way ANOVA with Dunnett’s multiple comparison test). Regions marked by dashed squares, expanded below with individual channels displayed (**a,i**, bottom). White arrowheads, co-localisation (**a,g,i**). Scale bars, 10 µm (**a,i**) and 2 µm (**g**).

Subsequently, the results obtained with CDC42 dominant-negative mutant were confirmed using the drug ML141, a specific inhibitor of CDC42 Rho GTPase activity. Cells treated with this compound showed a reduced CIP4 recruitment and actin polymerisation around all deformation sizes in a dose-dependent manner (**Figure 3d-e; Supplementary Figures 4d-e and 5d-e**). These observations suggest that CDC42 controls both local actin polymerisation and CIP4 recruitment. Interestingly, upon transient expression of each TOCA protein, we observed an increase in actin polymerisation around 100 nm deformations (**Figure 3g-h**) but no increase on 300 nm and 500 nm deformations (**Supplementary Figure 4g-h and 5g-h**).

Similarly, the knock-down of CIP4 slightly reduced actin polymerisation on 100 nm deformations (**Figure 3i-j**). This effect could be strongly enhanced by the concomitant depletion of all three TOCA family members, indicating that functional redundancy might exist between the three proteins (**Figure 3i-j**). Strikingly, no significant effect or only a slight reduction of actin polymerisation was observed upon CIP4 and all TOCA protein depletion around 300 nm and 500 nm deformations, respectively (**Supplementary Figures 4i-j and 5i-j**). Similarly, CDC42 depletion completely abolished the local actin polymerisation on all deformation sizes, while Rac1 or clathrin depletions, used here as negative controls, did not significantly decrease actin polymerisation on 300 nm and 500 nm deformations and led only to a partial decrease on 100 nm (**Figure 3i-j; Supplementary Figures 4i-j and 5i-j**). Accordingly, immunofluorescence experiments only showed a slight accumulation of clathrin around membrane deformations and no correlation with actin recruitment (**Supplementary Figure 7a-f**).

Taken together, these observations suggest that locally activated CDC42 recruits CIP4 (or other TOCA proteins) to plasma membrane nanodeformations, where they cooperate to induce local actin polymerisation. Strikingly, this CDC42-TOCA protein system seems to be highly curvature-sensitive, as it appears strictly required for local actin polymerisation on the smaller size deformations (100 nm), while not on larger sizes (300 nm and 500 nm). More precisely, while CDC42 seems to be important for actin ring formation around all deformation sizes, TOCAs are strictly required only on 100 nm deformations. This indicates that other partners may participate in controling actin polymerisation around larger membrane deformations. Moreover, these actin structures are not related to clathrin plaques as their formation seems independent of clathrin.

In order to identify the players involved in actin ring formation, we further investigated the regulation of CDC42, focusing on Rho GAP domain-containing proteins, since some of them are also known to be part of the BAR domain protein superfamily. Briefly, we individually depleted Nadrin1/2, Oligophrenin, srGAP1/2/3, and monitored actin polymerisation around 500 nm plasma membrane nanodeformations (**Supplementary Figure 8a-b**). Even though some of these proteins did not show any obvious enrichment around the deformations in the previous screening (see **Figure 2a**), the depletion of Nadrin2, Oligophrenin and srGAP3 significantly increased actin polymerisation (**Supplementary Figure 8a-b**). These observations suggest that these Rho GAP domain-containing BAR proteins are activators of CDC42 in the process studied here. Indeed, we could assume that upon their depletion, CDC42 remains activated, leading to an increase in actin polymerisation or to more stable actin structures.

Then, we drew our attention on the potential lipid species involved in our mechanism of curvature-mediated actin polymerisation. In particular, we explored the role of phosphoinositides (PIs) as they are well-known regulators of actin polymerisation at the plasma membrane [29]. In this context, PI(4,5)P_2_ is recognised as the major regulator thanks to its ability to recruit and activate several components of the actin polymerisation machinery at the plasma membrane, including N-WASP [30]. Moreover, it was shown that PI(4,5)P_2_ recruits CDC42 and FNBP1L to lipid bilayers where they promote Arp2/3-dependent actin polymerisation *in vitro* [13,15]. To test the involvement of PI(4,5)P_2_ in the current process, we used a rapalog-based system, allowing the controlled recruitment of a 5’-phosphatase to the cytosolic leaflet of the plasma membrane and subsequent depletion of PI(4,5)P_2_ [31]. Strikingly, upon induction of phosphatase recruitment to the cell surface, we observed a complete loss of local actin rings around all nanodeformation sizes, while other actin fibers seemed unaffected (**Supplementary Figure 8c-d**). Of note, the depletion of PI3P species by inhibiting PI3-kinases with wortmannin [32] showed no effect (or only mild effect) on actin polymerisation and CIP4 recruitment (**Supplementary Figure 8e-f**), suggesting that PI3Ps are not involved in the process. Altogether, these data indicate that curvature-mediated actin polymerisation strictly relies on the presence of PI(4,5)P_2_, but not PI(3,4,5)P_3_, in the cytosolic leaflet of the plasma membrane.

#### Local CIP4/CDC42-dependent actin polymerisation is dynamic and promoted by plasma membrane curvature

To validate that local CIP4 and CDC42-dependent actin polymerisation is promoted by plasma membrane nanoscale curvature, we performed live-cell microscopy experiments to measure protein enrichment in the vicinity of membrane nanodeformations over time. As previously described in a study using another strategy to generate nanostructured surfaces [12], we observed that the actin cytoskeleton (labeled with LifeAct) around plasma membrane deformations is dynamic, constantly undergoing rapid assembly and disassembly (**Movie 2**). Interestingly, pairwise observations of CDC42, CIP4 and actin showed that the three proteins follow very similar dynamics (**Figure 4a-c and Movies 3-5**). Of note, PalMyr-mCherry, used here as a plasma membrane marker, did not show any obvious dynamic behaviour (**Figure 4d and Movie 6**). To be more quantitative, we then computed the correlation coefficients between the dynamics of the three proteins, taken by pairs (**Figure 4e**). As expected, a positive and significant correlation was observed for the dynamics of CIP4/CDC42, CIP4/LifeAct and LifeAct/CDC42 couples, while no correlation was observed between the signals of PalMyr-mCherry and LifeAct-GFP, validating the involvement of CIP4 and CDC42 in curvature-induced actin polymerisation (**Figure 4e**). Interestingly, similar correlations were observed for the three proteins on 500 nm plasma membrane deformations (**Supplementary Figure 9a-e and Movies 7-9**). However, on these larger deformations, a significant correlation was also observed between the dynamics of actin and PalMyr-mCherry (**Supplementary Figure 9d-e and Movies 10**), suggesting that the distribution of the plasma membrane marker itself may be influenced by local actin polymerisation. This hypothesis is very likely, as actin polymerisation can promote the formation of lipid rafts in the plasma membrane and local enrichment in lipid-anchored proteins [33–36]. Alternatively, actin polymerisation may enhance membrane deformations and therefore increase PalMyr-GFP signal artificially.

**Figure 4:**
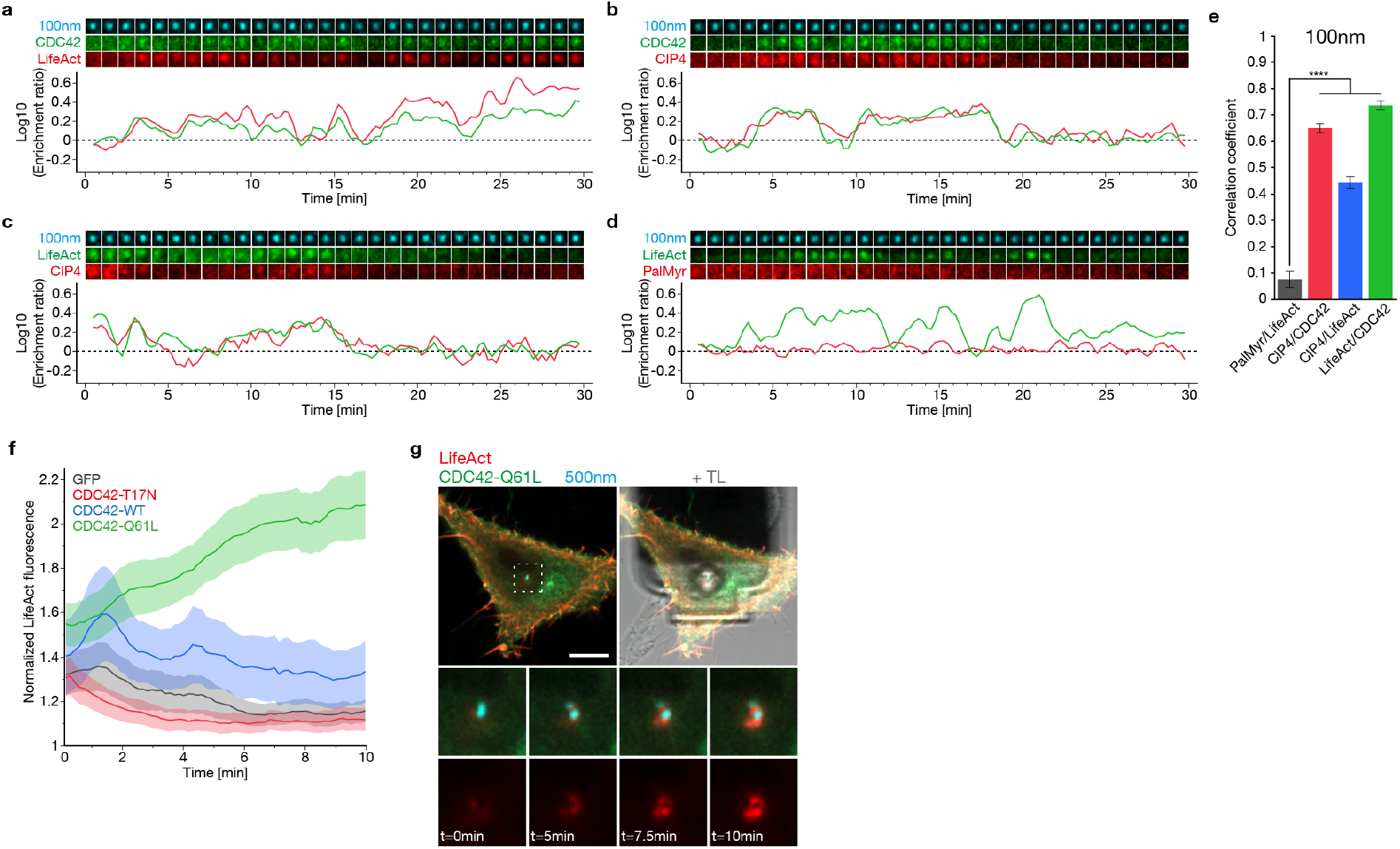
Local CIP4 and CDC42-dependent actin polymerisation is dynamic and promoted by plasma membrane curvature. **a-d**, Representative Airyscan time-lapses and the corresponding quantifications of enrichment ratios of the following pairs around 100 nm deformations over time: **a**, GFP-CDC42 and LifeAct-mCherry; **b**, GFP-CDC42 and mCherry-CIP4; **c**, LifeAct-GFP and mCherry-CIP4; **d**, LifeAct-GFP and PalMyr-mCherry (PalMyr). 30 min time-lapses with 15 sec intervals between frames. **e**, Correlation coefficients between the enrichment ratios over time curves for the pairs shown in **a-d**. Number of deformations: PalMyr/LifeAct, n = 65; CIP4/CDC42, n = 118; CIP4/LifeAct, n = 83; LifeAct/CDC42, n = 145. Three independent experiments. **f-g**, Monitoring of CDC42-dependent actin polymerisation upon deformation of the plasma membrane by FluidFM. **f**, Quantification of normalized LifeAct-mCherry fluorescence intensity around FluidFM tip over time upon co-expression of GFP (control, black), GFP-CDC42-T17N (red), GFP-CDC42-WT (blue) or GFP-CDC42-Q61L (green). 10 min time-lapses with 9 sec intervals between frames. Number of cells: GFP, n = 20; GFP-CDC42-T17N, n = 18; GFP-CDC42-WT, n = 15; GFP-CDC42-Q61L, n = 17. Two independent experiments. **g**, Representative image of FluidFM/confocal experiments upon co-expression of GFP-CDC42-Q61L and LifeAct-mCherry. Region marked by a dashed square surrounding the bead, expanded below at the indicated time points; t = 0 min, initial deformation of the cell membrane. TL, transmitted light (FluidFM cantilever) (representative images of GFP, GFP-CDC42-T17N and GFP-CDC42-WT in **Supplementary Figure 9f-h**). Data are mean ± s.e.m.; **** P < 0.0001 (**e**, one-way ANOVA with Dunnett’s multiple comparison test). Scale bar, 10 µm (**g**).

The deformation of plasma membrane using nanostructured cell culture substrates is a long process, which requires cell adhesion to proceed from 30 min to 2 hours. In order to grasp the dynamics of CDC42 and local actin polymerisation upon acute membrane deformation (within 10 min), we carried out additional experiments on live cells using a FluidFM (<u>Fluid</u>ic <u>F</u>orce <u>M</u>icroscopy) device coupled to a fast-scanning confocal microscope, already used in a previous study [37]. With this unique setup, we were able to induce plasma membrane nanodeformations in a time-, space- and force-controlled manner, while simultaneously imaging protein recruitment in the vicinity of the deformed plasma membrane. Fluorescent 500 nm particles were captured at the tip of the FluidFM probe and pulled towards the apical plasma membrane of cells co-expressing LifeAct-mCherry and different GFP-tagged CDC42 constructs. Interestingly, upon transient expression of wild-type CDC42, we observed a peak of actin polymerisation within 2 min after induction of deformation, followed by a progressive decrease of the signal (**Figure 4f, Supplementary Figure 9f and Movie 11**). A similar behaviour was observed in cells expressing free GFP, although the actin signal was lower overall (**Figure 4f, Supplementary Figure 9g and Movie 12**). Interestingly, expression of the dominant negative mutant of CDC42 (T17N) completely inhibited actin polymerisation around plasma membrane deformations (**Figure 4f, Supplementary Figure 9h and Movie 13**). By contrast, expression of the constitutively active CDC42 mutant (Q61L) led to a continuous increase in actin polymerisation throughout the 10 min duration of the experiment (**Figure 4f, Figure 4g and Movie 14**).

Taken together, these data show that plasma membrane nanodeformation promotes actin polymerisation in a CDC42-dependent manner, possibly through the direct activation of CDC42. Our data also show the highly dynamic nature of this mechanism *in cellulo*, and demonstrate that it can occur both at the basal and apical plasma membranes, despite their differences in composition and mechanical properties [38].

#### Plasma membrane nanodeformations can serve as tuning platforms for IFNγ-receptor signaling

It is well reported in literature that the cortical actin cytoskeleton is an essential regulator of plasma membrane protein organisation and function [34,39]. Given the drastic local reorganisation of actin observed upon growing cells on nanostructured substrates, one may wonder whether this reorganisation could affect signaling pathways. Here, we investigated the signaling of the IFNγ-R, which is one of the major signaling pathways in cells known to be involved in tumour immune-surveillance, tumour escape, but also in a broad range of inflammatory diseases such as cardiovascular disease [40,41]. In the current model, IFNγ-R is a complex composed of 4 chains: two chains IFNγ-R1 and two chains IFNγ-R2. When IFNγ binds to the IFNγ-R complex, a conformational change of the chains occurs leading to the activation of pre-associated Janus-<u>a</u>ctivated tyrosine <u>k</u>inases (JAK), which triggers the phosphorylation of <u>s</u>ignal transducer and <u>a</u>ctivator of transcription 1 (STAT1) resulting in its nuclear translocation [42]. Importantly, IFNγ-R is known to be regulated by actin - and lipid – nanodomains as the signaling of a mutant of IFNγ-R displaying additional glycosylations was shown to be strongly inhibited due to its trapping in the actin meshwork caused by galectin 1 or 3 binding [43].

First, the subcellular localisation of IFNγ-R2 subunit was monitored in HeLa cells expressing IFNγ-R2-GFP seeded on 500 nm nanostructures. Interestingly, the enrichment in IFNγ-R2-GFP around membrane deformations was systematically correlated with actin enrichment (**Figure 5a**, white arrowheads). If actin was not enriched, there was no enrichment in IFNγ-R2-GFP (**Figure 5a**, purple arrowhead). To be more physiological and avoid potential artefacts due to transient expression, we used Caco-2 cells, which, on the contrary to Hela cells, express IFNγ-R endogenously and respond to IFNγ stimulation. IFNγ-R2 signal was monitored by immunofluorescence after 10 min of stimulation with IFNγ. Although the signal was more discrete/punctated, we observed endogenous IFNγ-R2 around 500 nm deformations (**Figure 5b**, yellow arrowheads).

**Figure 5:**
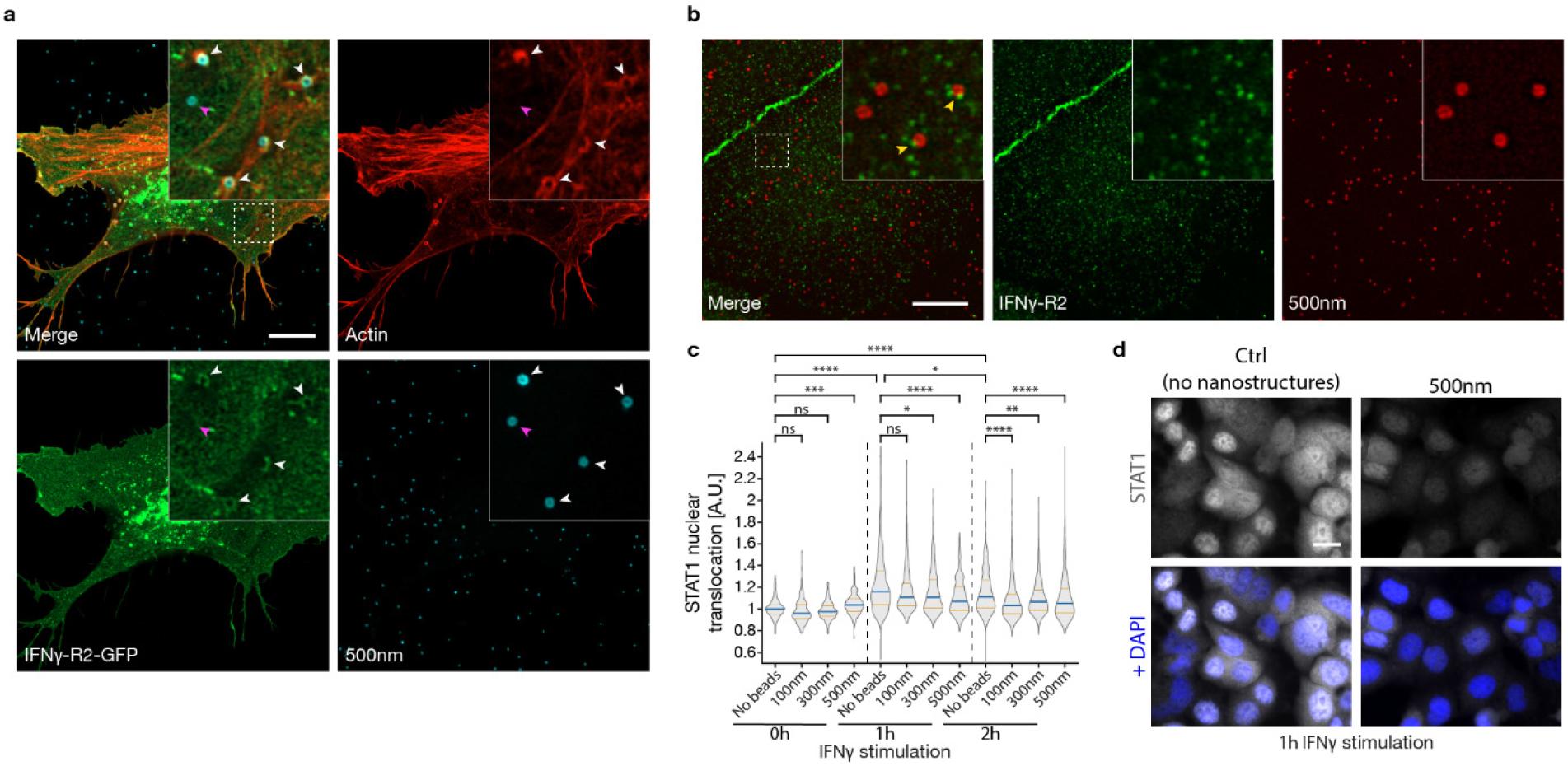
Plasma membrane nanodeformations can serve as platforms for IFNγ-receptor signaling regulation. HeLa (**a**) or Caco-2 (**b-d**) cells. **a**, Transient expression of IFNγ-R2-GFP (green) without stimulation in cells grown on 500 nm nanostructures (blue). Red signal, actin (phalloidin). Note the co-distribution of IFNγ-R2-GFP and actin signals (white arrowheads), or the simultaneous absence of both signals (purple arrowhead) around nanodeformations. **b**, Endogenous IFNγ-R2 (green) after 10 min of stimulation with IFNγ in cells grown on 500 nm nanostructures (red). Note the punctated localisation of IFNγ-R2 around nanodeformations (yellow arrowheads). **a-b**, Representative Airyscan images of 3 independent experiments. Insets, zoom on dashed square regions. **c-d**, STAT1 nuclear translocation experiment upon IFNγ stimulation for indicated times in cells grown on nanostructured substrates. Quantifications shown in **c**. Representative images after 1h stimulation (**d**; representative images of all conditions in **Supplementary Figure 10**). Number of cells: No beads/0h, n = 308; 100 nm/0h, n = 242; 300 nm/0h, n = 365; 500 nm/0h, n = 347; No beads/1h, n = 499; 100 nm/1h, n = 401; 300 nm/1h, n = 507; 500 nm/1h, n = 494; No beads/2h, n = 548; 100 nm/2h, n = 560; 300 nm/2h, n = 502; 500 nm/2h, n = 701. Data are median with quartiles from two (100 nm) or three (No beads, 300 nm and 500 nm) independent experiments. **** P < 0.0001; *** P < 0.001; ** P < 0.01; * P < 0.05 (Kruskal–Wallis test with Dunn’s multiple comparison test). Scale bars, 10 µm (**a-b**) and 20 µm (**d**).

Consequently, the next logical step was to investigate whether the presence of IFNγ-R in the curvature-induced actin nanodomains could affect its downstream signaling. Therefore, STAT1 nuclear translocation was monitored in Caco-2 cells seeded for 1.5 hour on flat or nanostructured substrates, and then treated for the indicated times with IFNγ before fixation (**Figure 5c-d and Supplementary Figure 10**). In the absence of IFNγ stimulation, STAT1 nuclear translocation did not occur and stayed at a basal level in all conditions, except for 500 nm deformations where a slightly higher basal level was observed (**Figure 5c**, 0h). As expected, STAT1 nuclear translocation increased after IFNγ stimulation in cells grown on surfaces without nanostructures, with a peak at 1 hour (**Figure 5c-d**, 1h and 2h). Strikingly, STAT1 nuclear translocation was significantly lower when cells were grown on nanostructures (**Figure 5c-d**). Furthermore, increasing nanostructure size led to a stronger inhibition of IFNγ-STAT1 signaling pathway, with a maximal effect reached with 500 nm nanostructures after 1 hour of treatment with IFNγ (**Figure 5c-d**).

Altogether, these results highlight that IFNγ-R is localised in actin nanodomains formed around plasma membrane deformations and that its downstream IFNγ-induced JAK/STAT1 signaling is severely impaired by the presence of the deformations induced by the nanostructures.

## Discussion

Our screening dissociated the two functions of BAR domain proteins - as membrane curvature sensors vs inducers - to solely study their ability to sense/bind to pre-existing deformations *in vivo*. We found that BAR proteins from all subfamilies - excepted PX-BAR - were able to bind membrane deformations of every tested size. Hence, there seems to be no apparent correlation between the intrinsic curvature radius of BAR proteins and their affinity for a particular deformation size. Especially, both N-BAR and F-BAR proteins - usually thought to bind highly or shallowly curved membrane, respectively - were recruited on all deformation sizes. Thus, the classical view suggesting that BAR domain protein subfamily classification depicts their ability to bind/induce distinct membrane curvature ranges is challenged by our results [18–20]. Their curvature range affinity seems more protein specific than family-/intrinsic curvature radius-dependent.

Furthermore, multiple proteins bound to a range of curvature radii while others were restricted to a more specific curvature radius. A very likely explanation for this observation resides in the flexibility and plasticity of some BAR oligomer scaffolds to adapt to broad membrane curvature ranges as shown for CIP4 *in silico* [44]. Our screening results showed that for each size of deformation, a unique set of BAR domain proteins bound to it. These unique sets of BAR proteins would allow cells to adapt and react in a specific way depending on the nanogeometry of their plasma membrane. Different cellular processes could be primed at these sites of high membrane curvature, which could be of importance for mechanotransduction. In our model, TOCA proteins are necessary for actin polymerisation to occur only on 100 nm membrane deformations. Other BAR domain proteins could fulfil their role regarding actin polymerisation on bigger deformations. In particular, SNX9 was shown to be able to replace FNBP1L and to recruit N-WASP to mediate actin polymerisation when PI3P is present in curved supported lipid bilayers instead of PI(4,5)P_2_ [15]. Yet, SNX9 was only recruited at a low level to membrane deformations in our screening. Unlike other BAR domain proteins, its recruitment might be short-lived or restricted to only a subpopulation of membrane deformations. Therefore, additional research to observe the dynamics of SNX9 and actin cytoskeleton on membrane deformations is needed to assess the potential involvement of this protein in the current mechanism.

While the TOCA subfamily has been previously involved in curvature-mediated actin polymerisation [12], our study is the first to connect the Rho GTPase CDC42 to this mechanism *in cellulo*. Specifically, our results suggest that the activity of CDC42, an early actor involved in actin polymerisation, might be curvature-dependent. Although we did not provide any direct demonstration, several indirect observations in our study argue in favour of this hypothesis. We observed changes in curvature-mediated actin polymerisation by modulating CDC42 activity using constitutively active and dominant negative mutants, inhibitors and more importantly by the transient expression of Rho GAP proteins which are well-known to target CDC42. Indeed, we showed that some Rho GAP-containing BAR domain proteins - Nadrin2, Oligophrenin and srGAP3 - decreased actin polymerisation around plasma membrane nanodeformations, most likely by deactivating CDC42. This could possibly explain the transient nature of actin rings. However, we have not yet identified Rho <u>g</u>uanine nucleotide-<u>e</u>xchange factors (GEF) candidates which could activate CDC42 in the first place. Surprisingly, there are very few Rho GEF proteins among the BAR superfamily. Although very challenging, additional experiments will thus be necessary to monitor directly CDC42 activity, to establish if it is regulated by membrane curvature itself and to identify the GEFs involved in the mechanism.

In addition, our RNA interference experiments showed that CDC42 - and not Rac1 - is necessary for curvature-mediated actin polymerisation to occur, which was previously only demonstrated in *in vitro* studies on liposomes, and flat or curved supported lipid bilayers [13,15]. Remarkably, the observations made by Gallop et al. [15] using an *in vitro* model system share multiple similarities with our observations, such as the involvement of CDC42, FNBP1L and PI(4,5)P_2_ in local actin polymerisation. After activation of CDC42 and recruitment of TOCAs to membrane deformations, it seems likely that actin polymerisation occurs through N-WASP and Arp2/3 complex activation [12]. These aspects will require further investigations in our experimental setup.

Strikingly, the plasma membrane nanodomains described in our study are reminiscent of early-stage FEME-like clathrin-independent endocytic pits. In FEME, locally activated CDC42 recruits CIP4 and FBP17 to prime the membrane for endocytosis before being inactivated thanks to Rho GAP-containing BAR domain proteins (Nadrin1, SH3BP1 and Oligophrenin) in the absence of receptor activation or stabilisation [21]. Most of these various characteristic molecular actors are involved in the membrane nanodomains we described here. This opens up the exciting possibility of the existence of a frustrated clathrin-independent endocytosis, as has been shown with the frustrated clathrin-mediated endocytosis previously described [22,23]. While further research is required to support this hypothesis, these frustrated structures could also serve as signaling platforms promoting the accumulation of receptors, adaptor proteins or acting as a scaffold for the organisation of the actin cytoskeleton.

In particular, previous studies have already reported the formation of actin structures on topography-induced membrane deformations, connecting them to various cellular functions such as adhesion, migration or endocytosis [10,12,45,46]. Here, we propose that these structures can also act as platforms regulating cell surface receptor signaling. We showed that IFNγ-R colocalises with actin rings on membrane deformations and that its signaling is significantly impaired by the presence of nanostructures. Interestingly, Blouin et al. [43] showed previously that IFNγ-R signaling was inhibited when the receptor is trapped in the actin meshwork. We propose that a similar regulation occurs on topography-induced plasma membrane deformations, which enhance the formation of numerous and dense actin structures (*i*.*e*. actin rings) throughout the membrane. This could increase the probability for IFNγ-R of being trapped, preventing its signaling. Additionally, it is tempting to establish a direct conceptual analogy between the FEME-like structures described in our study and frustrated clathrin-mediated endocytosis described by other groups, where clathrin plaques were shown to act as platforms regulating receptor signaling (in particular for epidermal growth factor receptor) [22–24]. Yet, additional investigations are necessary to characterise IFNγ-R dynamics within these topography-induced actin nanodomains. It will also be of great interest to explore if other receptors are trapped in these actin structures and whether their signaling is also affected.

In conclusion, our study sheds light on the poorly explored connection between plasma membrane nanogeometry and signal transduction mechanisms. Importantly, we identified an additional mechanotransduction pathway induced by substrate nanotopography, distinct from clathrin plaques. This pathway involves the BAR domain protein CIP4, the Rho GTPase CDC42 and actin cytoskeleton, and is capable of tuning type II interferon signaling. These findings may be key to understand how substrate nanotopography influences cell behaviour in biological contexts as diverse as development, cancer progression or for the design of biocompatible materials, to cite only a few examples.

## Materials and Methods

### Antibodies and other reagents

The following antibodies were purchased from the indicated suppliers: mouse monoclonal anti-CIP4 (Santa Cruz Biotechnology, sc-135868, 1:500 for immunofluorescence and 1:1000 for Western blot); mouse monoclonal anti-CDC42 (Cytoskeleton Inc., ACD03, 1:250 for Western blot); mouse monoclonal anti-Rac1 (Merck Millipore, 05-389, 1:1000 for Western blot); mouse monoclonal anti-clathrin heavy chain (BD Biosciences, 610500, 1:100 for immunofluorescence and 1:5000 for Western blot); mouse monoclonal anti-α-tubulin (Sigma, T5168, 1:5000 for Western blot); mouse monoclonal anti-β-actin (Santa Cruz Biotechnology, sc-47778, 1:1000 for Western blot); rabbit polyclonal anti-IFNγ-R2 (proteintech, 10266-1-AP, 1:200 for immunofluorescence); rabbit monoclonal anti-STAT1 (D1K9Y) (Cell Signalling Technology, 14994, 1:300 for immunofluorescence); secondary antibodies conjugated to Alexa Fluor 488 or 647 (Thermo Fisher Scientific, 1:250 for immunofluorescence); anti-mouse secondary antibody conjugated to STAR ORANGE (Abberior, STORANGE-1001, 1:200 for immunofluorescence); anti-mouse and anti-rabbit secondary antibodies conjugated to horseradish peroxidase (Sigma and Dako, respectively, 1:5000 for Western blot). The rabbit polyclonal anti-FBP17 (1:1000 for Western blot) was a kind gift from Pietro De Camilli (New Haven, USA) and the mouse monoclonal anti-FNBP1L (1:25 for Western blot) was a kind gift from Emanuela Frittoli (Milan, Italy). Phalloidin conjugated to Alexa Fluor 488 (A12379) or Alexa Fluor 633 (A22284), Alexa Fluor 488 NHS ester (A20000), Alexa Fluor 555 NHS ester (A20009), rapamycin (PHZ1235) and PageRuler™ Prestained Protein Ladder (26617) were purchased from Thermo Fisher Scientific. Phalloidin conjugated to STAR RED (STRED-0100) was purchased from Abberior. ATTO390 (AD 390) and ATTO647N (AD 647N) were purchased from ATTO-TEC. Phalloidin conjugated to iFluor 555 (ab176756) was purchased from Abcam. Poly(allylamine) (PAH) solution (479144) and ML141 (SML0407) were purchased from Sigma. Silica colloids of 500 (24323-15), 300 (24321-15) and 100 nm (24041-10) were purchased from Polysciences. Wortmannin (HY-10197) was purchased from MedChemExpress. Human IFNγ was purchased from Imukin.

#### Synthesis of nanostructured substrates

##### Fluorescent silica colloids synthesis

Spherical silica colloids of 500 nm (0.075% wt.) were first coated with 0.025% wt. PAH solution for 30 min at room temperature. They were then washed 6 times by centrifugation at 10.000 xg followed by resuspension in water. Next, PAH-coated colloids were fluorescently labeled with 10 μg m^-1^ NHS-ester conjugated dyes (ATTO-390, Alexa Fluor 488, Alexa Fluor 555 or ATTO-647N) for 1h at room temperature in water. Colloids were washed 6 more times by centrifugation at 21.000 xg followed by resuspension in water. The fluorescent colloid suspension was then stored at 4°C protected from any light until further use. For colloids of 300 and 100 nm, the procedure was the same except they were coated with 0.05% wt. PAH, and colloids of 100 nm were labeled with 20 μg ml^-1^ of fluorescent dyes.

##### Fluorescent silica colloids adsorption

Fluorescent silica colloids were diluted in water to 0.024% wt. for 500 nm, 0.0045% wt. for 300 nm and 0.0006% wt. for 100 nm and transferred to a 10 mm optical glass fluorescent cuvette. Two coverslips (WillCo Wells, 12 mm in diameter) were pressed together, placed vertically in the cuvette and incubated for 2 hours in the dark at room temperature. After incubation, water was slowly added at the top of the cuvette and the same volume was removed at the bottom. This washing procedure was repeated 6 times, with the last 3 being carried out with 100% isopropanol (VWR). Coverslips were then separated and individually dipped in isopropanol before being placed in 4 well plates, nanostructures facing up. Coverslips were left to dry and kept in the dark at room temperature until further use.

For live cell imaging, nanostructures were created on coverslips of 35 mm in diameter in the same way and assembled into dishes using WillCo-dish®KIT “Do-It-Yourself”.

##### Collagen coating

Dry nanostructured coverslips were incubated with 100 μg ml^-1^ collagen (Bornstein and Traub Type I, Sigma Aldrich, C8919) in PBS (Sigma Aldrich) for 1 hour in the dark at room temperature. Coverslips were then washed twice with PBS and twice with ethanol 70% (VWR). The collagen-coated coverslips were kept up to 48 hours before cell seeding.

##### Surface characterisation

Collagen coated nanostructured substrates were sputter coated with gold before imaging with a JEOL 7600F scanning electron microscope.

#### Cell culture

HeLa (human cervix adenocarcinoma, ATCC® CCL-2) were grown at 37 °C under 5% CO_2_ in DMEM high glucose glutamax (Gibco, 61965-059) supplemented with 10% FBS, 100 U ml^−1^ penicillin, 100 µg ml^−1^ streptomycin, and 1 mM pyruvate.

Caco-2 (colorectal adenocarcinoma, ATCC® HTB-37) were grown at 37 °C under 5% CO_2_ in DMEM low glucose pyruvate (Gibco, 31885-023) supplemented with 10% FBS, 100 U ml^−1^ penicillin and 100 µg ml^−1^ streptomycin.

#### DNA constructs and transfection

For all DNA transfection experiments except for IFNγ-R2-eGFP, plasmids were transfected with FuGene HD (Promega, E2311) in HeLa cells according to the manufacturer’s instructions. Cells were used for experiments 16-24 h after transfection. IFNγ-R2-eGFP plasmid was transfected in HeLa or Caco-2 cells with X-tremeGENE HP DNA transfection reagent (Roche, 6366236001) according to the manufacturer’s instructions, for 24h and with the following ratio: 1 µg DNA per 2 µL transfection reagent. All the plasmid used can be found in **Table 1** and **Table 2**.

**Table 1:** Plasmids containing BAR domain protein sequences used in this study.

| Protein name | Other names | Species | Backbone | Source | Additional information |
| --- | --- | --- | --- | --- | --- |
| Arfaptin2 |  | Human | pCI | Harvey T. McMahon (Cambridge, UK) | BAR domain only |
| FAM92B |  | Human | pCI | Harvey T. McMahon (Cambridge, UK) |  |
| ICA1L |  | Human | pCI | Harvey T. McMahon (Cambridge, UK) | BAR domain only |
| ICA69 |  | Human | pCI | Harvey T. McMahon (Cambridge, UK) | BAR domain only |
| PICK1 |  | Rat | pEYFP-C1 | Madsen K.L. et al. [47] |  |
| Amphiphysin | AMPH | Human | pEGFP-C1 | Yamada H. et al. [48] |  |
| Bin1 | Amphiphysin II | Human | pEGFP-C1 | Lee E. et al. [49] |  |
| Bin2 |  | Human | pEGFP-N3 | Maria Jose Sanchez-Barrena (Madrid, Spain) |  |
| Bin3 |  | Mouse | pEGFP-N1 | François Tyckaert (Louvain-la-Neuve, Belgium) |  |
| Endophilin A1 | SH3GL2 | Human | pcDNA3 | Harvey T. McMahon (Cambridge, UK) |  |
| Endophilin A2 | SH3GL1 | Rat | pEGFP-N1 | Pietro De Camilli (New Haven, USA) |  |
| Endophilin A3 | SH3GL3 | Human | pcDNA3 | Harvey T. McMahon (Cambridge, UK) |  |
| Endophilin B1 |  | Human | pEYFP-N1 | Karbowski M. et al. [50] |  |
| Endophilin B2 |  | Human | pCI | Harvey T. McMahon (Cambridge, UK) | BAR domain only |
| Nadrin1 | RICH1, ARHGAP17 | Human | pEYFP-Nx | Isabell Begemann (Münster, Germany) |  |
| Nadrin2 | RICH2, ARHGAP44 | Human | pEYFP-Nx | Isabell Begemann (Münster, Germany) | BAR domain only |
| SH3BP1 |  | Human | pmCherry-Cx | Maria Carla Parrini (Paris, France) |  |
| ACAP1 | Centaurin B1 | Human | pEGFP-C1 | Li J. et al. [51] |  |
| ACAP2 | Centaurin B2 | Mouse | pEGFP-C1 | Mitsunori Fukuda (Sendai, Japan) |  |
| ACAP3 | Centaurin B5 | Human | pCI | Harvey T. McMahon (Cambridge, UK) | BAR domain only |
| APPL1 |  | Mouse | pmCherry-C1 | Addgene #27683 |  |
| ASAP2 |  | Human | pCI | Harvey T. McMahon (Cambridge, UK) | BAR domain only |
| Graf1 | ARHGAP26 | Human | pEGFP-C3 | Richard Lundmark (Umeå, Sweden) |  |
| Graf2 | ARHGAP10 | Human | pCI | Harvey T. McMahon (Cambridge, UK) | BAR domain only |
| Oligophrenin1 |  | Human | pCAMCS | Linda Van Aelst (New York, USA) |  |
| SNX1 |  | Human | pEGFP-C1 | Pete Cullen (Bristol, UK) |  |
| SNX2 |  | Human | pEGFP-C1 | Pete Cullen (Bristol, UK) |  |
| SNX4 |  | Human | pEGFP-C1 | Pete Cullen (Bristol, UK) |  |
| SNX5 |  | Human | pEGFP-C1 | Paul Gleeson (Melbourne, Australia) |  |
| SNX6 |  | Human | pEGFP-C1 | Pete Cullen (Bristol, UK) |  |
| SNX7 |  | Human | pEGFP-C1 | Pete Cullen (Bristol, UK) |  |
| SNX8 |  | Human | pEGFP-C1 | Pete Cullen (Bristol, UK) |  |
| SNX9 |  | Mouse | pmCherry-C1 | Addgene #27678 |  |
| SNX18 |  | Human | pDEST | Anne Simonsen (Oslo, Norway) |  |
| SNX30 |  | Human | pEGFP-C1 | Pete Cullen (Bristol, UK) |  |
| SNX32 |  | Human | pEGFP-C2 | Pete Cullen (Bristol, UK) |  |
| SNX33 |  | Human | pEGFP-C2 | Pete Cullen (Bristol, UK) |  |
| ARHGAP4 |  | Human | pEGFP-C1 | Kathy Gould (Nashville, USA) |  |
| ARHGAP4 |  | Human | pEGFP-C1 | Kathy Gould (Nashville, USA) | BAR domain only |
| ARHGAP29 |  | Human | pcDNA3 | Johannes Bos (Utrecht, The Netherlands) |  |
| CIP4 | TRIP10, TOCA2 | Mouse | pmCherry-C1 | Addgene #27685 |  |
| FBP17 | TOCA3 | Human | pmCherry-C1 | Addgene #27688 |  |
| FCHo1 |  | Mouse | pmCherry-C1 | Addgene #27690 |  |
| FCHo2 |  | Mouse | pmCherry-C1 | Addgene #27686 |  |
| FCHSD1 |  | Mouse | pEGFP-C2 | Zhigang Xu (Shandong, China) |  |
| FCHSD2 |  | Mouse | pEGFP-C2 | Zhigang Xu (Shandong, China) |  |
| Fer | TYK3 | Human | pEGFP-C1 | Kathy Gould (Nashville, USA) |  |
| Fer | TYK3 | Human | pEGFP-C1 | Kathy Gould (Nashville, USA) | BAR domain only |
| Fes | Fps | Human | pEGFP-C1 | Kathy Gould (Nashville, USA) | BAR domain only |
| FNBP1L | TOCA1 | Human | pEGFP-C1 | Pietro De Camilli (New Haven, USA) |  |
| GAS7 |  | Human | pEGFP-N1 | Chuck C.K. Chao (Taoyuan, Taiwan) |  |
| GMIP |  | Mouse | pEGFP-N1 | Kazunobu Sawamoto (Nagoya, Japan) |  |
| HMHA1 | ARHGAP45 | Human | pEGFP-C3 | Peter Hordijk (Amsterdam, The Netherlands) |  |
| Nostrin |  | Human | pEGFP-C1 | Kathy Gould (Nashville, USA) | BAR domain only |
| PACSIN1 | Syndapin1 | Mouse | pEGFP-N3 | Shiro Suetsugu (Ikoma, Japan) |  |
| PACSIN2 | Syndapin2 | Mouse | pmCherry-C1 | Addgene #27681 |  |
| PACSIN3 | Syndapin3 | Mouse | pEGFP-N3 | Shiro Suetsugu (Ikoma, Japan) |  |
| PSTPIP1 |  | Human | pEGFP-C2 | Deborah Gumucio (Ann Arbor, USA) |  |
| PSTPIP2 |  | Human | pCI | Harvey T. McMahon (Cambridge, UK) |  |
| srGAP1 | ARHGAP13 | Human | pCAG-EGFP | Franck Polleux (New York, USA) |  |
| srGAP1 | ARHGAP13 | Human | pCAG-EGFP | Franck Polleux (New York, USA) | BAR domain only |
| srGAP2 | ARHGAP34 | Human | pCAG-EGFP | Franck Polleux (New York, USA) |  |
| srGAP2 | ARHGAP34 | Mouse | pCAG-mRFP | Franck Polleux (New York, USA) | BAR domain only |
| srGAP3 | ARHGAP14 | Human | pCAG-EGFP | Franck Polleux (New York, USA) |  |
| srGAP3 | ARHGAP14 | Human | pCAG-EGFP | Franck Polleux (New York, USA) | BAR domain only |
| IRSp53 | BAIAP2 | Human | pCI | Harvey T. McMahon (Cambridge, UK) | BAR domain only |
| IRTKS | BAIAP2L1 | Human | pCI | Harvey T. McMahon (Cambridge, UK) | BAR domain only |
| MIM |  | Human | pCI | Harvey T. McMahon (Cambridge, UK) | BAR domain only |
| Pinkbar | BAIAP2L2 | Human | pCI | Harvey T. McMahon (Cambridge, UK) | BAR domain only |

**Table 2:** Other plasmids used in this study.

| Protein name | Species | Backbone | Source |
| --- | --- | --- | --- |
| PalMyr-GFP | / | pEGFP-C1 | Daniel Gehrlich (Vienna, Austria) |
| PalMyr-mCherry | / | pmCherry-C1 | Gaëlle Boncompain (Paris, France) |
| LifeAct-GFP | / | pIRES | Philippe Chavrier (Paris, France) |
| LifeAct-mCherry | / | pIRES | Philippe Chavrier (Paris, France) |
| GFP | / | pEGFP-C1 | Clontech (discontinued) |
| CDC42-WT | Mouse | pEGFP-C1 | Anne Ridley (London, UK) |
| CDC42-T17N | Human | pEGFP-C1 | Florence Niedergang (Paris, France) |
| CDC42-Q61L | Human | pcDNA3 | Addgene #12600 |
| Rac1-WT | Human | pEGFP-C1 | Florence Niedergang (Paris, France) |
| PM-FRB-GFP | / | pEGFP-N1 | [31] provided by Volker Haucke (Berlin, Germany) |
| mRFP-FKBP12-5'Ptase | / | pmRFP-C1 | [31] provided by Volker Haucke (Berlin, Germany) |
| IFNγ-R2 | Human | pcDNA3.1 V5/His | Cédric Blouin (Paris, France) |

#### RNA interference

siRNAs used in this study were purchased from Qiagen and transfected with HiPerFect (Qiagen, 301704) according to the manufacturer’s instructions. Experiments were always performed 72 h after siRNA transfection, where protein depletion efficiency was maximal as shown by immunoblotting analysis with specific antibodies. AllStars Negative Control siRNA (Qiagen, 1027281) served as a reference point. Depletion of each protein was achieved using 1 to 4 different sequences (**Table 3**) used as a pool at a total final concentration of 40 nM. To deplete CIP4, FBP17 and FNBP1L, two siRNA per protein were used at a total final concentration of 40 nM.

**Table 3:**
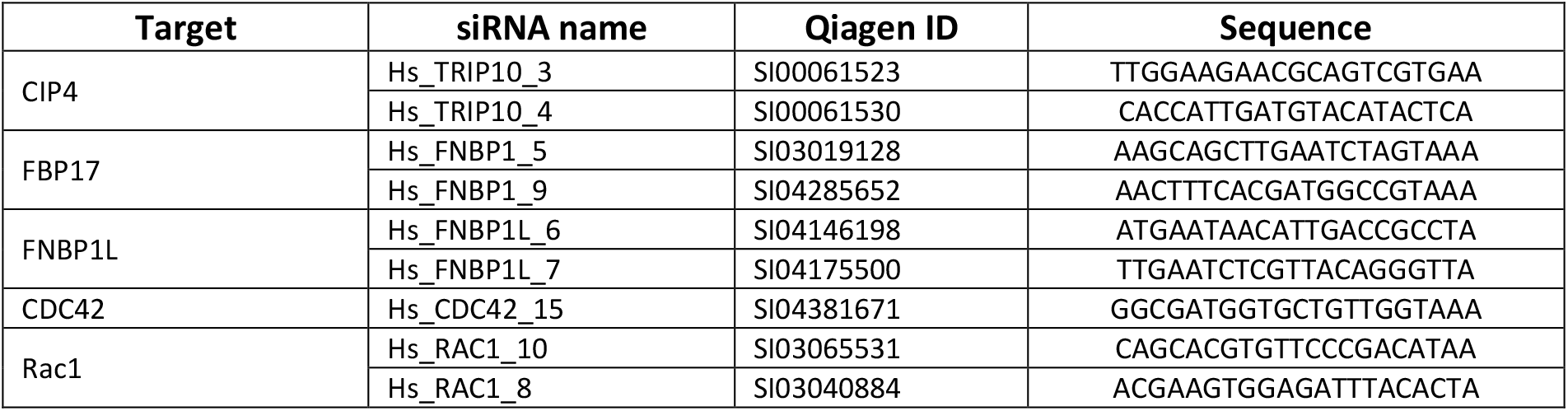
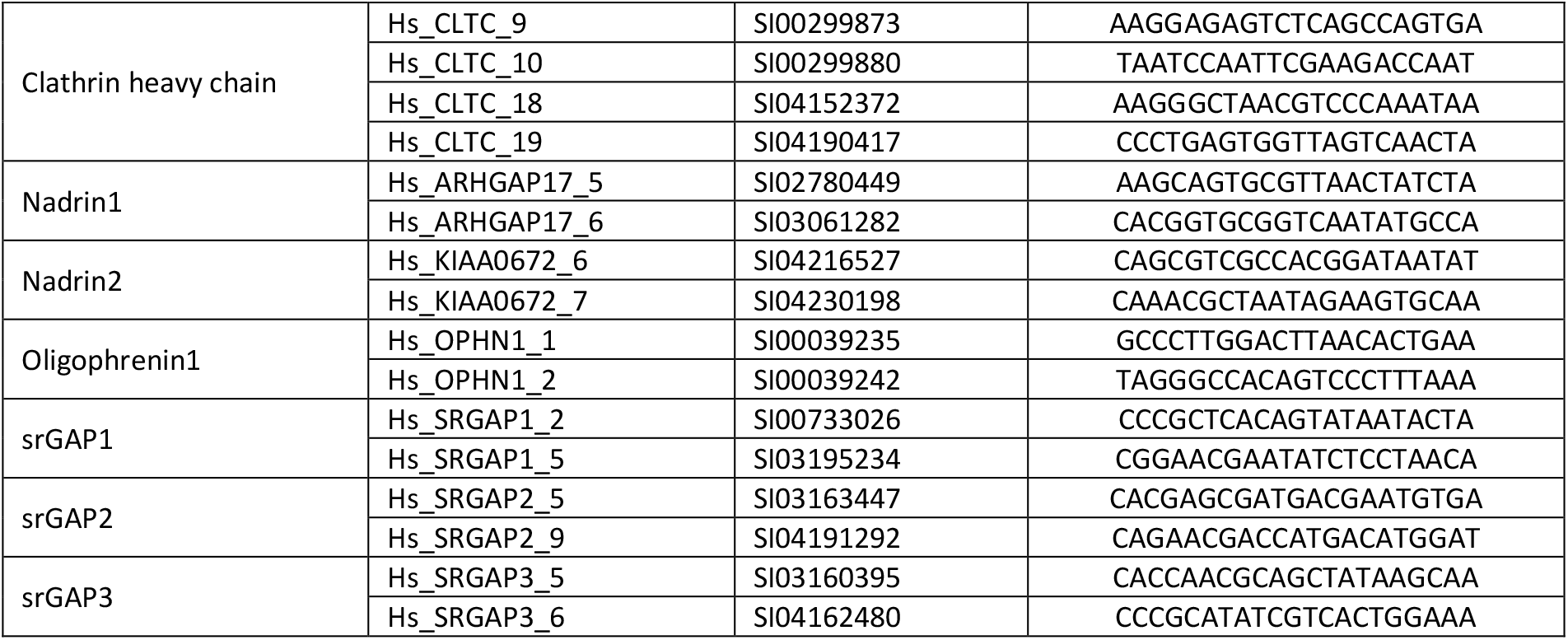
siRNAs used in this study.

#### Dose-dependent inhibition of CDC42

Cells were pre-incubated for 1 hour in medium containing 1 % DMSO (Sigma) (control condition), 20 µM, 50 µM or 100 µM ML141 in DMSO. Cells were then harvested and seeded on collagen coated nanostructured substrates under the same ML141 concentrations for 1 hour before fixation and labeling.

#### Phosphoinositide level manipulation

For the depletion of PI3 species, cells were seeded on collagen coated nanostructured substrates and either 1 µM Wortmannin in DMSO or 0.1% DMSO (control condition) was immediately added to the medium. Cells were cultured for 1 hour on the substrates before fixation and labeling.

For the rapalog-based system used to deplete PI(4,5)P_2_, cells previously transfected with both PM-FRB-GFP and mRFP-FKBP12-5’Ptase were seeded on collagen coated nanostructured substrates and cultured for 50 min. Next, 20 nM rapamycin in DMSO or 0.2% DMSO (control condition) was added to the medium for 10 min before cells fixation and labeling.

#### STAT1 nuclear translocation assay

Caco-2 cells were seeded at 80% confluency for 1.5 hours in complete medium on collagen coated nanostructured substrates. Cells were gently washed twice with PBS and then starved for 20 min in medium without serum, to avoid any signaling background. Next, cells were stimulated or not with IFNγ 1000 U ml^−1^for 1 or 2 hours before fixation and labeling.

#### Immunofluorescence

For all experiments except STAT1 nuclear translocation assay, cells were fixed with 4% paraformaldehyde (PFA) (Electron Microscopy Sciences, 15710) at 37 °C for 15 min before quenching with 50 mM NH_4_Cl for at least 15 min. If needed, cells were then permeabilized with saponin (0.02% saponin (Sigma, 47036), 0.2% bovine serum albumin (Sigma, A9647) in PBS) for 30 min. Cells were then incubated with primary and secondary antibodies and/or phalloidin for 50 min in saponin, and mounted in Fluoromount G (Invitrogen).

For STAT1 nuclear translocation assay, cells were fixed with 4% PFA for 10 min at room temperature, quenched in 50 mM NH_4_Cl for 10 min and permeabilized with 0.1% Triton-X100 in 0.2% BSA in PBS for 10 min. After blocking with 0.2% BSA in PBS for 20 min, cells were incubated with primary anti-STAT1 antibody for 45 min at room temperature followed by secondary antibody labeling. To finish, coverslips were mounted in Fluoromount G (eBioscience) with 2 µg ml^−1^ DAPI (Sigma) to label nuclei.

#### Light microscopy

Fixed samples were imaged with a 34-channel Zeiss LSM 710 or LSM 900 confocal microscope equipped with an Airyscan detector and with a Plan Apo 63× numerical aperture (NA) 1.4 oil immersion objective. Wide-field images were acquired on a Zeiss LSM 900 microscope equipped with a sCMOS Prime 95B camera and a Plan Apo 40× numerical aperture (NA) 1.3 oil immersion objective.

2D STED super-resolution imaging was performed with a STEDYCON (Abberior Instruments GmbH) mounted on an upright microscope base (Zeiss AxioImager Z.2) equipped with a Zeiss Plan Apochromat 100x, NA 1.4 oil immersion objective actuated by a z-piezo with 100 microns range (Physik Instruments). In STED mode, 100 % of a max. nominal STED laser output of 1.25 W (775 nm) was used, leading to a theoretical spatial resolution of 35-40 nm. STAR RED and STAR ORANGE were excited by pulsed laser sources at 640 nm and 561 nm respectively. Pinhole size was 1.13 Airy units at 650 nm. Gated detection in single photon counting mode was performed using Avalanche photodetectors.

For live-cell imaging, the previously described Zeiss LSM microscopes were equipped with stage-top incubation chambers, allowing temperature and CO_2_ control. Cells were imaged 30 min after seeding on nanostructured substrates. Several time series of 30 min were then recorded over the course of 2 hours. Observations were made at 37 °C and 5% CO_2_.

#### Correlative FluidFM/fast-scanning confocal microscopy

For the dynamic study of actin recruitment upon defined membrane deformation, we used a combined instrument, made up of a FluidFM setup (Cytosurge) coupled to a Bruker Resolve AFM and to a Zeiss LSM980 fast-scanning confocal microscope equipped with a Plan Apo 63× numerical aperture (NA) 1.4 water immersion objective.

HeLa cells previously seeded in 40 mm glass-bottom dishes (WillCo-dish) were transfected with the required plasmids 16-24 h before the experiment. Culture media were refreshed just before starting the experiment. Observations were made at 37 °C and 5% CO_2_.

A suspension of fluorescent silica colloids labeled with ATTO-647N was placed on a glass-bottom petri dish and imaged with confocal microscopy. A FluidFM microchanneled cantilever was approached to the surface of the dish and used to trap individual nanoparticle, by applying a steady negative pressure of -800 mbar through the microfluidic system. Using the force control of the Bruker Resolve atomic force microscope, the cantilever with the trapped nanoparticle was then approached to the membrane of individual transfected HeLa cells by applying a 5 nN force that was kept constant after the tip was engaged. In parallel, the focal plane of the confocal microscope was focused on the bead, and 10 min time series were recorded in fast-scanning mode to monitor fluorescence variations in the surroundings of the nanoparticle.

#### Image analyses and quantifications

Images were processed with Zen blue 3.3 (Zeiss) and Fiji 1.53f51 (NIH), all image quantifications were performed with Fiji. A macro was written to semi-automate fluorescence quantification around each individual nanostructure (see Supplementary Figure 2 for a representation of the quantification process). The macro is available upon reasonable request.

For Airyscan images of cells seeded on nanostructured substrates, regions of interest (ROIs) were drawn around cells and a threshold was manually applied to isolate the fluorescence of nanostructures from the background in these ROIs. Next, individual nanostructures were isolated based on their area and circularity using the plugin “Extended Particle Analyzer”. The plugin returns the coordinates of the centres of each nanostructure which were then overlaid on the channel to quantify. When a small drift between channels was observed, centres were realigned accordingly. Then, circular radial profiles were computed in the channels to quantify based on the overlaid nanostructure centres using the plugin “Radial Profile Angle”. All individual nanostructures were always analysed except for nanostructures that were too close to cell edges, which were excluded from the quantification. These radial profiles represent the average fluorescence intensity at a given distance from the centre. Each profile is normalized by the fluorescence intensity at 1 µm from the centre of the deformation, which correspond to the non-deformed/flat plasma membrane (0.5 µm for 100 nm nanostructures). The graph obtained represents the average increase in fluorescence intensity (called enrichment ratio) at a given distance from the centre of the nanostructure compared to the flat membrane. Enrichment ratios for each nanostructure were then pooled together and the average radial intensity profile was calculated. The average intensity at 250, 150 and 50 nm from the centre, for 500, 300 and 100 nm nanostructures respectively, was then used to compare various conditions.

For Airyscan time series of cells seeded on nanostructured substrates, stacks were first corrected for slight drifts during imaging using the plugin “HyperStackReg”. Of note, no correction for fluorophore bleaching was made as the enrichment ratios later computed are normalized on each frame. The plugin “Z project” was then used to average the fluorescence intensity of the nanostructures in a single image to refine their signals and to precisely find their centres. The quantification was then done similarly as previously described using the average centres to compute the profiles for each frame around each nanostructure. Quantification was done only on a subset of nanostructures. Nanostructures too close to cell edges were excluded and a fluctuation of either CDC42, CIP4, LifeAct or PalMyr signal around membrane deformations was required for their quantifications. Rings around membrane deformations that were stables for the entire duration of the acquisition were not quantified as there is no dynamic to extract. Extracted enrichment ratios over time for each channel were processed by first computing the unweighted moving averages and then correlation coefficients between the two signals for each deformation using JMP Pro v16.0 software.

For confocal time series acquired during FluidFM/confocal experiments, a circular ROI was defined around the fluorescent bead and the average fluorescence intensity values from the LifeAct channel was extracted from each frame. This average intensity was normalized by the LifeAct signal in a region away from the bead.

Nuclear translocation of STAT1 was quantified with a homemade plugin [52] by calculating the nucleo-cytoplasmic ratio of phospho-STAT1 signal. Nuclei masks were realized thanks to DAPI staining.

### Statistical analyses

All statistical analyses were performed using JMP Pro v16.0 software. To normalize the enrichment ratios into Gaussian distributions, a log_10_ transformation was systematically applied to the ratios before averaging them. In the case of Gaussian distributions, parametric tests were used, and data were represented on graphs as mean ± s.e.m. as error bars. In the case of non-Gaussian distributions, non-parametric tests were used, and data were represented on graphs as median with quartiles. Details on the parametric and non-parametric tests used for each analysis, as well as other statistical details related to specific graphs, are indicated in figure legends. Significance of comparisons is represented on the graphs by asterisks. No statistical method was used to predetermine sample size.

For the screening of BAR domain proteins on 500, 300 and 100 nm deformations (**Figure 2a-c**), independent experimental repeats were weighted in such a way that each one is equally represented in the pooled dataset.

## Acknowledgements

We would like to acknowledge the following people for help in experiments and providing materials or expertise: I. Begemann, C. Blouin, G. Boncompain, J. Bos, C.C.K. Chao, P. Chavrier, P. Cullen, P. De Camilli, M. Fukuda, D. Gehrlich, P. Gleeson, K. Gould, D. Gumucio, V. Haucke, P. Hordijk, R. Lundmark, H.T. McMahon, F. Niedergang, M.C. Parrini, F. Polleux, A. Ridley, M.J. Sanchez-Barrena, K. Sawamoto, Simonsen, S. Suetsugu, F. Tyckaert, D. Tyteca, L. Van Aelst and Z. Xu. We thank B. Knoops, P. Dumont, and F. Gofflot from LIBST (UCLouvain) for sharing cell culture facility and materials. We greatly acknowledge IMABIOL and Morph-Im imaging platforms from UCLouvain and UNamur respectively, and M.-C. Eloy for technical assistance. We thank the MICA platform from UCLouvain and D. Magnin for SEM imaging. We thank SMCS platform from UCLouvain and C. Rasse for advices regarding statistical analyses. We thank Abberior Instruments GmbH for allowing us to acquire STED images of our samples on their demonstration device. We thank A. Forrester for proofreading the manuscript. B.L. is supported by a PhD fellowship from FRIA/FNRS (Belgium). D.A. is a Research Associate from the FNRS. P.M. is supported by grants from the “Fonds National de la Recherche Scientifique” (FNRS, CDR-J.0119.19) and the “Communauté française de Belgique–Actions de Recherches Concertées” (17/22-085). H.-F.R. is supported by a Start-Up Grant Collen-Francqui from Francqui Foundation and an Incentive Grant for Scientific Research from the “Fonds de la Recherche Scientifique” (FNRS, MIS-F.4540.21).

