## Supplementary Information for "Plasma membrane nanodeformations promote actin polymerisation through CIP4/CDC42 recruitment and regulate type II IFN signaling"

**Ledoux *et al.***

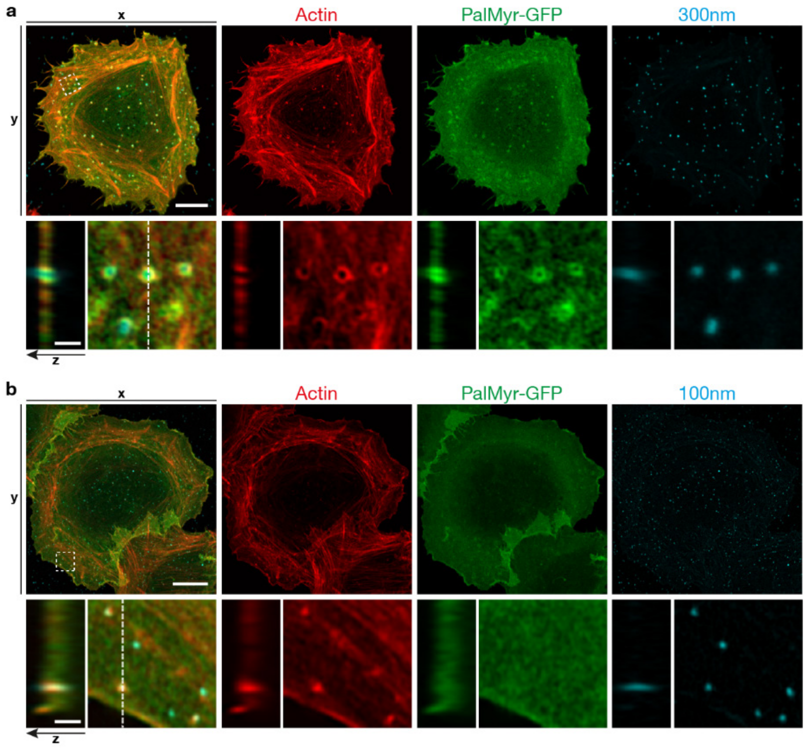


**Supplementary Figure 1: Nanostructures deform cell plasma membrane and induce local actin polymerisation.** HeLa cell expressing PalMyr-GFP (green) seeded on 300 nm (**a**) or 100 nm (**b**) nanostructures (cyan) and stained for actin (phalloidin, red). Regions marked by dashed squares, expanded below (panel **a-b**, bottom) and also displayed as transversal views along Z-axis. Scale bars, 10 µm (**a-b**, top) and 1 µm (**a-b**, bottom).

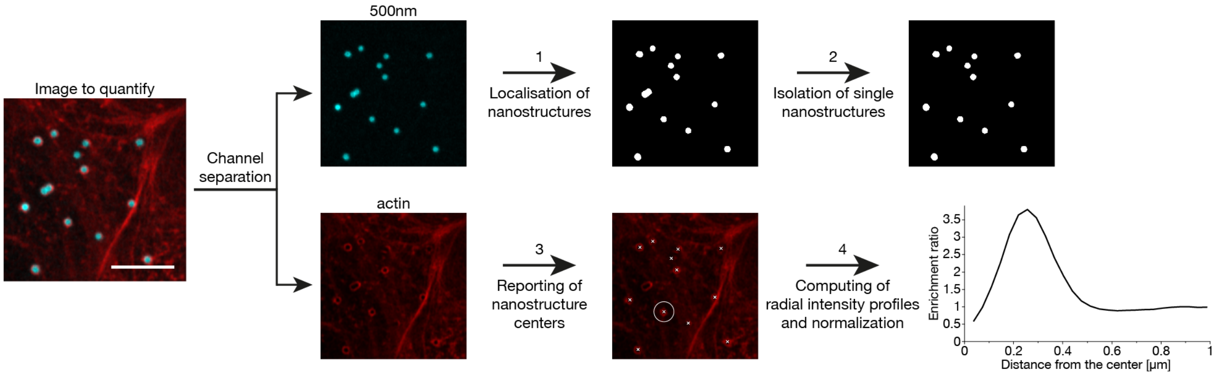


**Supplementary Figure 2: Workflow of fluorescence quantification around nanostructures.** First, nanostructures are found using a threshold (**1**) and sorted based on their size and shape to remove aggregates (**2**). Then, nanostructure centres are computed and reported in the channel to be quantified in order to create circular ROIs (**3**). Finally, local fluorescence intensity is quantified using a circular radial profile and normalized by the fluorescence intensity at 1 µm away from the centre of the deformation, which correspond to the non-deformed/flat membrane (0.5 µm for 100 nm nanostructures) (**4**). The graph obtained represents the average increase in fluorescence intensity (called enrichment ratio) at a given distance from the centre of the nanostructure compared to the flat membrane. Normalized intensities for each nanostructure are then pooled together and the average radial intensity profile is calculated. The average intensity at 250 nm, 150 nm and 50 nm away from the centre - for 500, 300 and 100 nm nanostructures, respectively - is then used to compare various conditions. Scale bar, 5 µm.

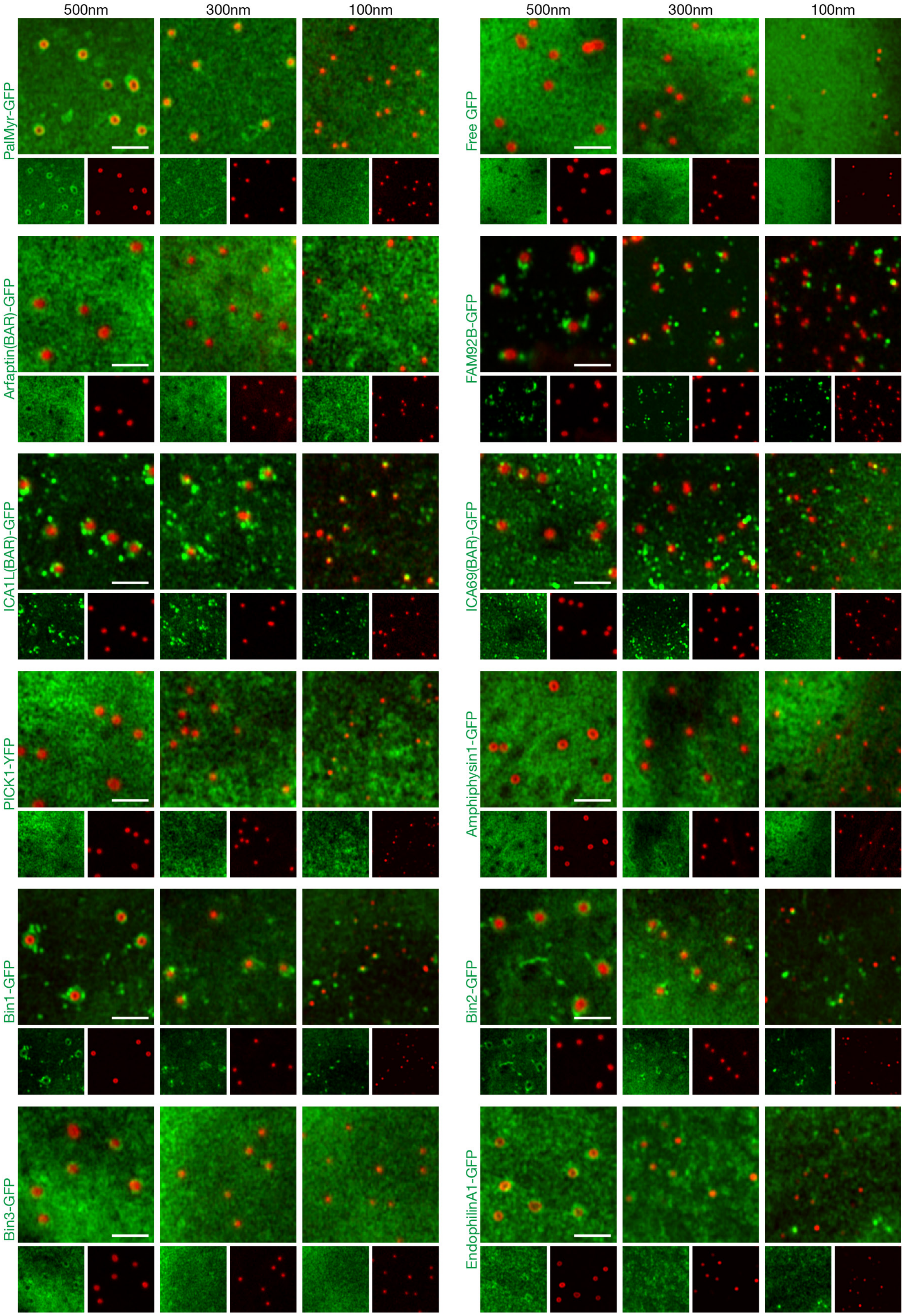


**Supplementary Figure 3: Representative images of all BAR domain proteins screened for their affinity to 500 nm, 300 nm and 100 nm plasma membrane deformations (part 1/6).** Scale bars, 2 µm.

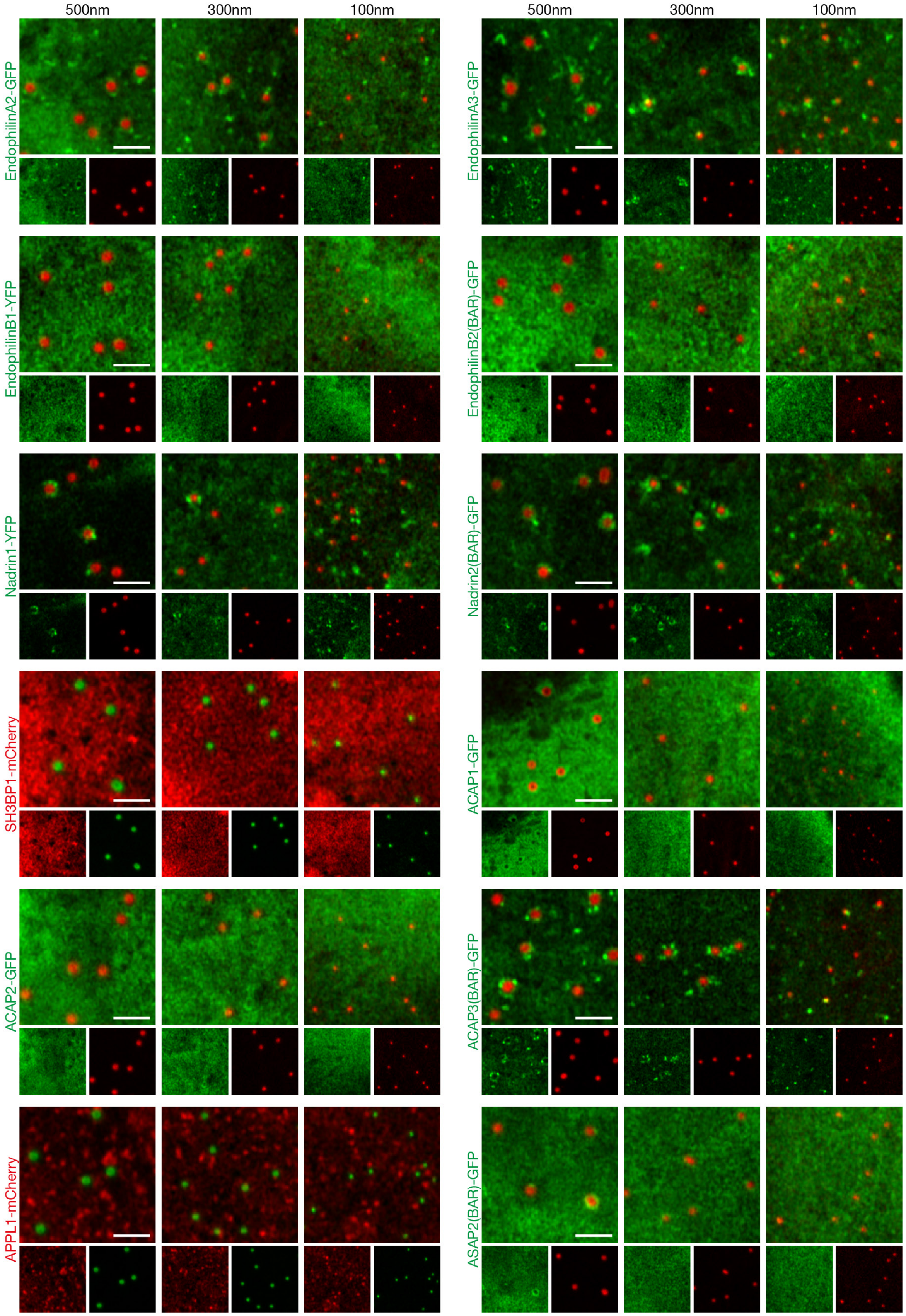


**Supplementary Figure 3: Representative images of all BAR domain proteins screened for their affinity to 500 nm, 300 nm and 100 nm plasma membrane deformations (part 2/6).**

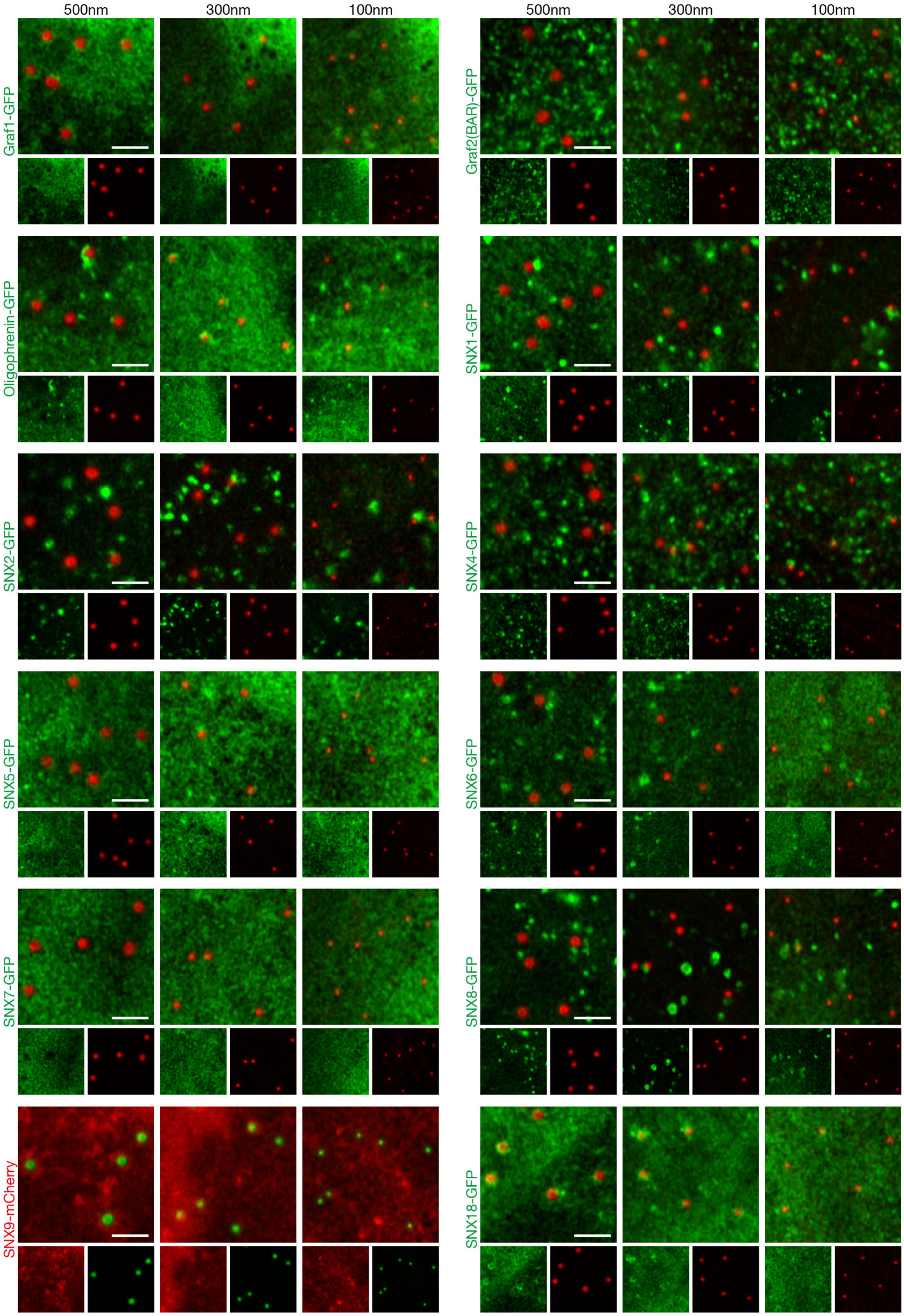


**Supplementary Figure 3: Representative images of all BAR domain proteins screened for their affinity to 500 nm, 300 nm and 100 nm plasma membrane deformations (part 3/6).**

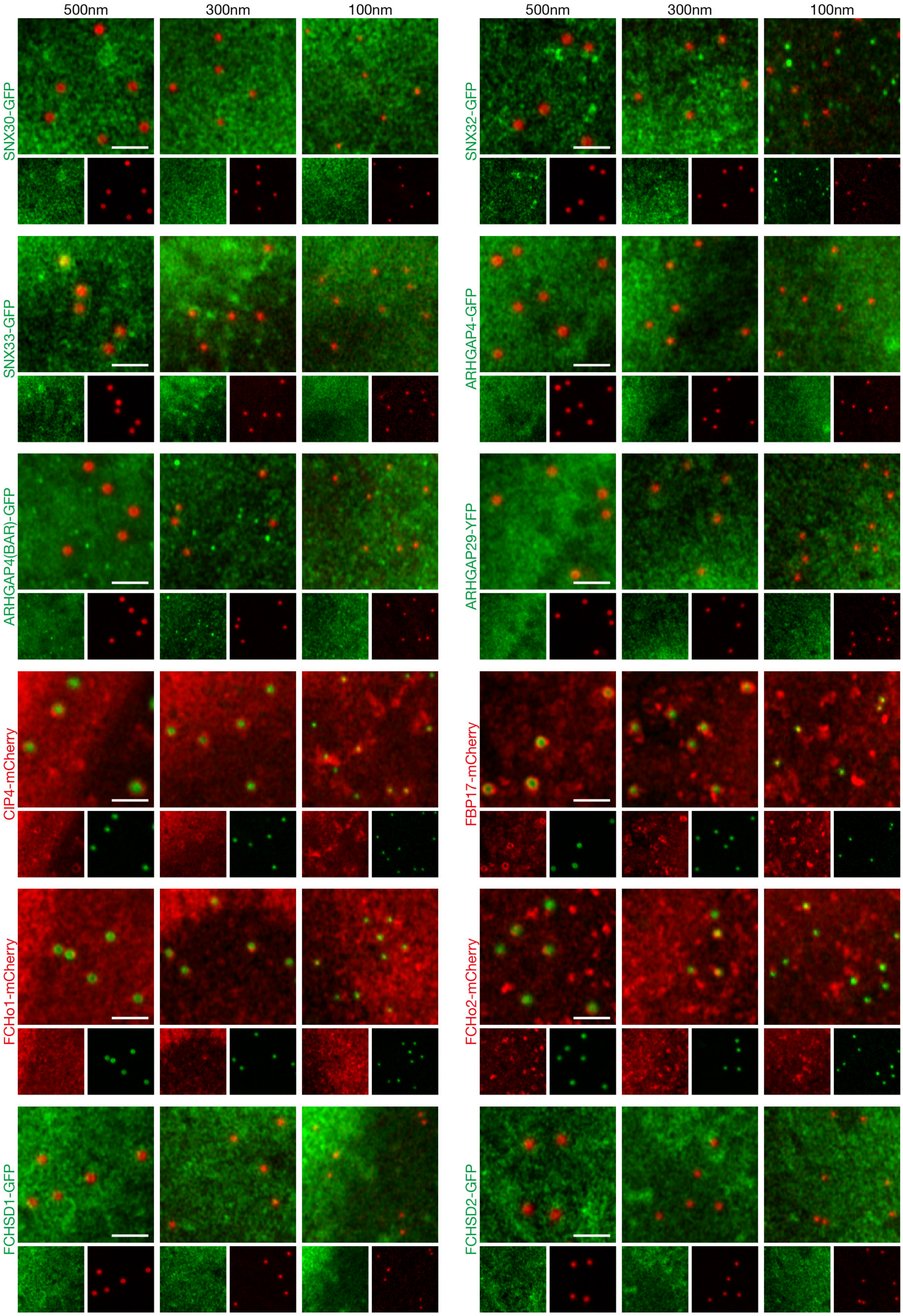


**Supplementary Figure 3: Representative images of all BAR domain proteins screened for their affinity to 500 nm, 300 nm and 100 nm plasma membrane deformations (part 4/6).**

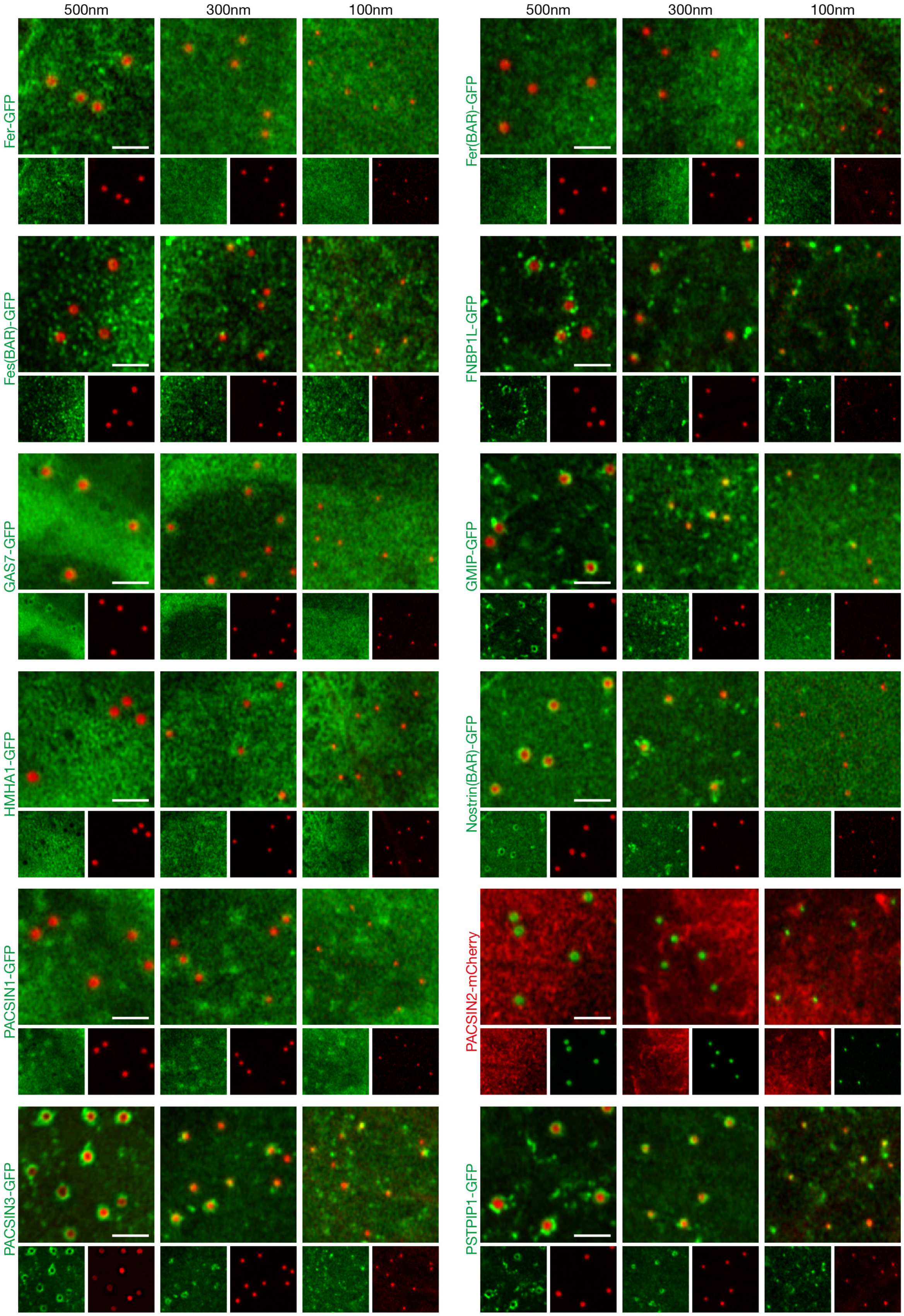


**Supplementary Figure 3: Representative images of all BAR domain proteins screened for their affinity to 500 nm, 300 nm and 100 nm plasma membrane deformations (part 5/6).**

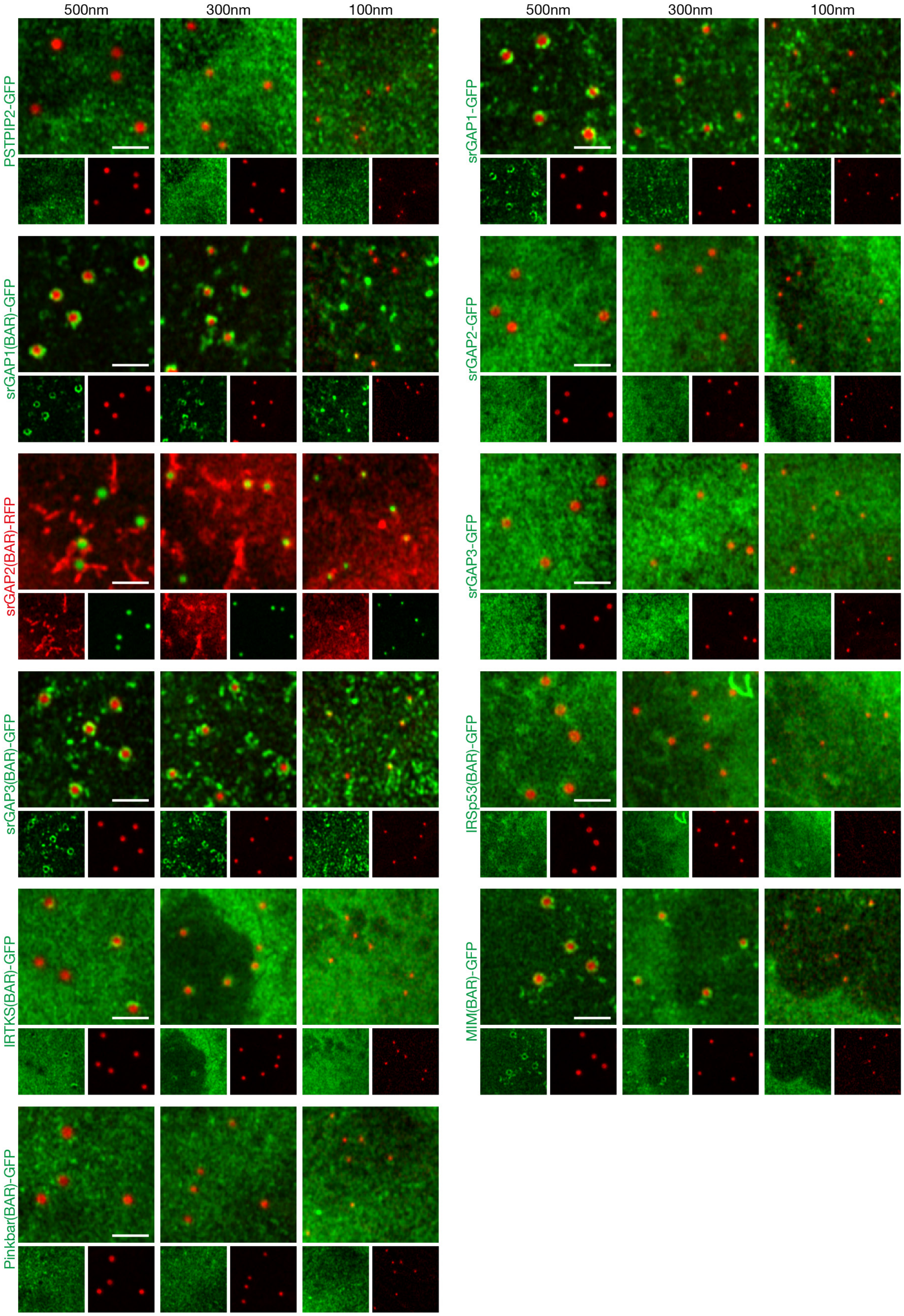


**Supplementary Figure 3: Representative images of all BAR domain proteins screened for their affinity to 500 nm, 300 nm and 100 nm plasma membrane deformations (part 6/6).**

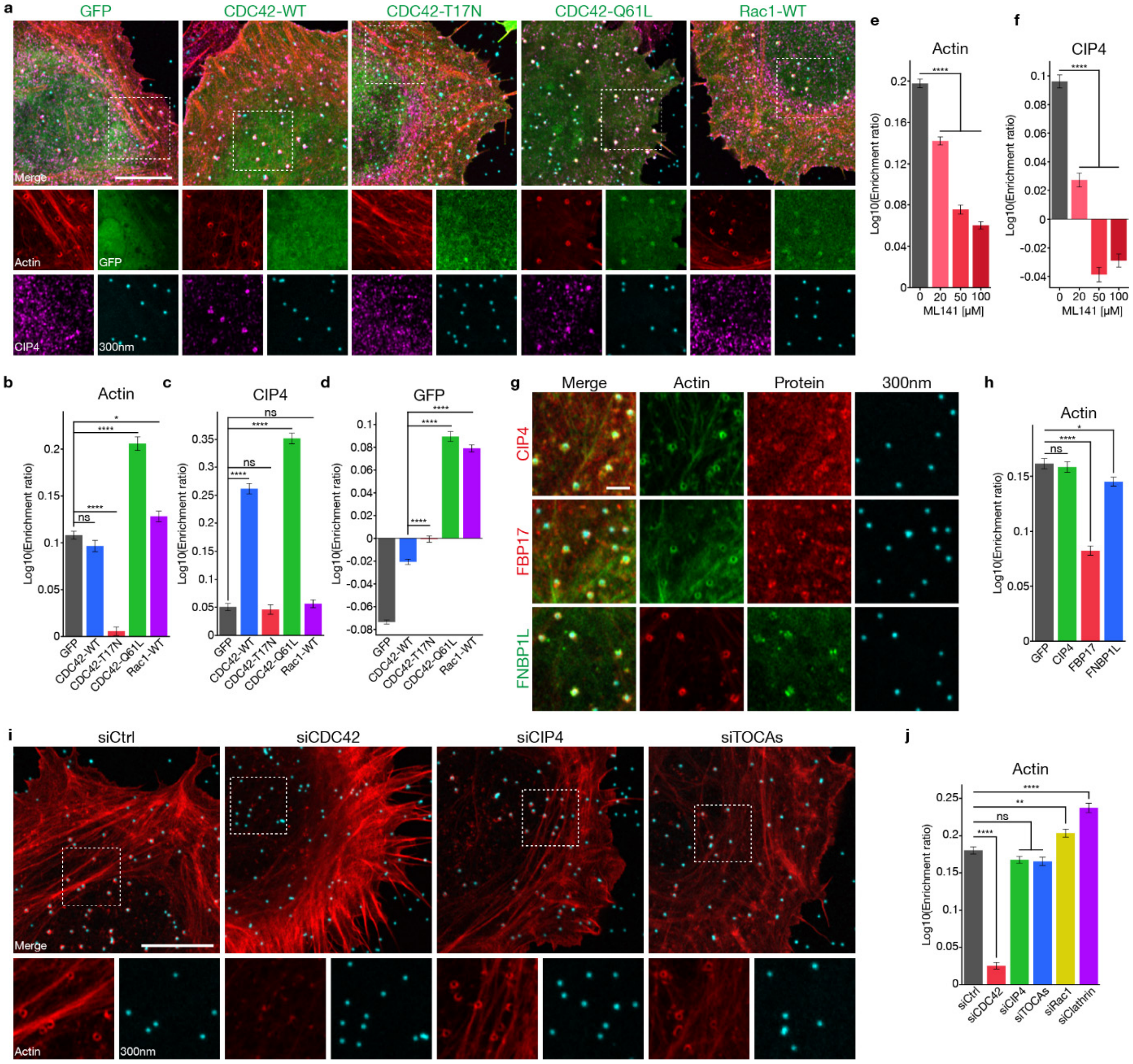


**Supplementary Figure 4: CDC42 and CIP4 control actin polymerisation around 300 nm plasma membrane deformations.** HeLa cells grown on 300 nm nanostructures and transfected with fluorescent constructs (**a-d,g-h**), treated with a specific CDC42 inhibitor (ML141) (**e-f**) or with siRNAs (**i-j**), as indicated. Data shown are quantifications of actin (**b,e,h,j**), endogenous CIP4 (**c,f**) or GFP (**d**) fluorescence around 300 nm deformations and the corresponding representative Airyscan images (**a,g,i**), as indicated. **a-d**, Effect of transient expression of GFP, GFP-CDC42-WT, GFP-CDC42-T17N, GFP-CDC42-Q61L or GFP-Rac1-WT, on the enrichment of actin, CIP4 and GFP around deformations. Number of deformations: GFP, n = 1880; CDC42-WT, n = 1587; CDC42-T17N, n = 1156; CDC42-Q61L, n = 1399; Rac1-WT, n = 1574. Three independent experiments. **e-f**, Effect of dose-dependent inhibition of CDC42 by ML141 (0 µM, 20 µM, 50 µM or 100 µM) on the enrichment of actin and CIP4 around deformations. Number of deformations: 0 µM, n = 2704; 20 µM, n = 2464; 50 µM, n = 1900; 100 µM, n = 2389. Three independent experiments. **g-h**, Effect of transient expression of GFP or individual TOCA proteins (mCherry-CIP4, mCherry-FBP17 or GFP-FNBP1L) on the enrichment of actin around deformations. Number of deformations: GFP, n = 2018; CIP4, n = 2362; FBP17, n = 2064; FNBP1L, n = 2748. Four independent experiments. **i-j**, Effect of the depletion of CDC42, CIP4, TOCAs (CIP4, FBP17 and FNBP1L), Rac1 or Clathrin Heavy Chain with siRNAs on the enrichment of actin around deformations. Number of deformations: siCtrl, n = 2002; siCDC42, n = 1062; siCIP4, n = 1960; siTOCA, n = 1797; siRac1, n = 2057; siClathrin, n = 1459. Three independent experiments. Data are mean ± s.e.m.; ns, not significant; **** $P < 0.0001$; ** $P < 0.01$; * $P < 0.05$ (one-way ANOVA with Dunnett's multiple comparison test). Regions marked by dashed squares, expanded below with individual channels displayed (**a,i**, bottom). White arrowheads, co-localisation (**a,g,i**). Scale bars, 10 µm (**a,i**) and 2 µm (**g**).

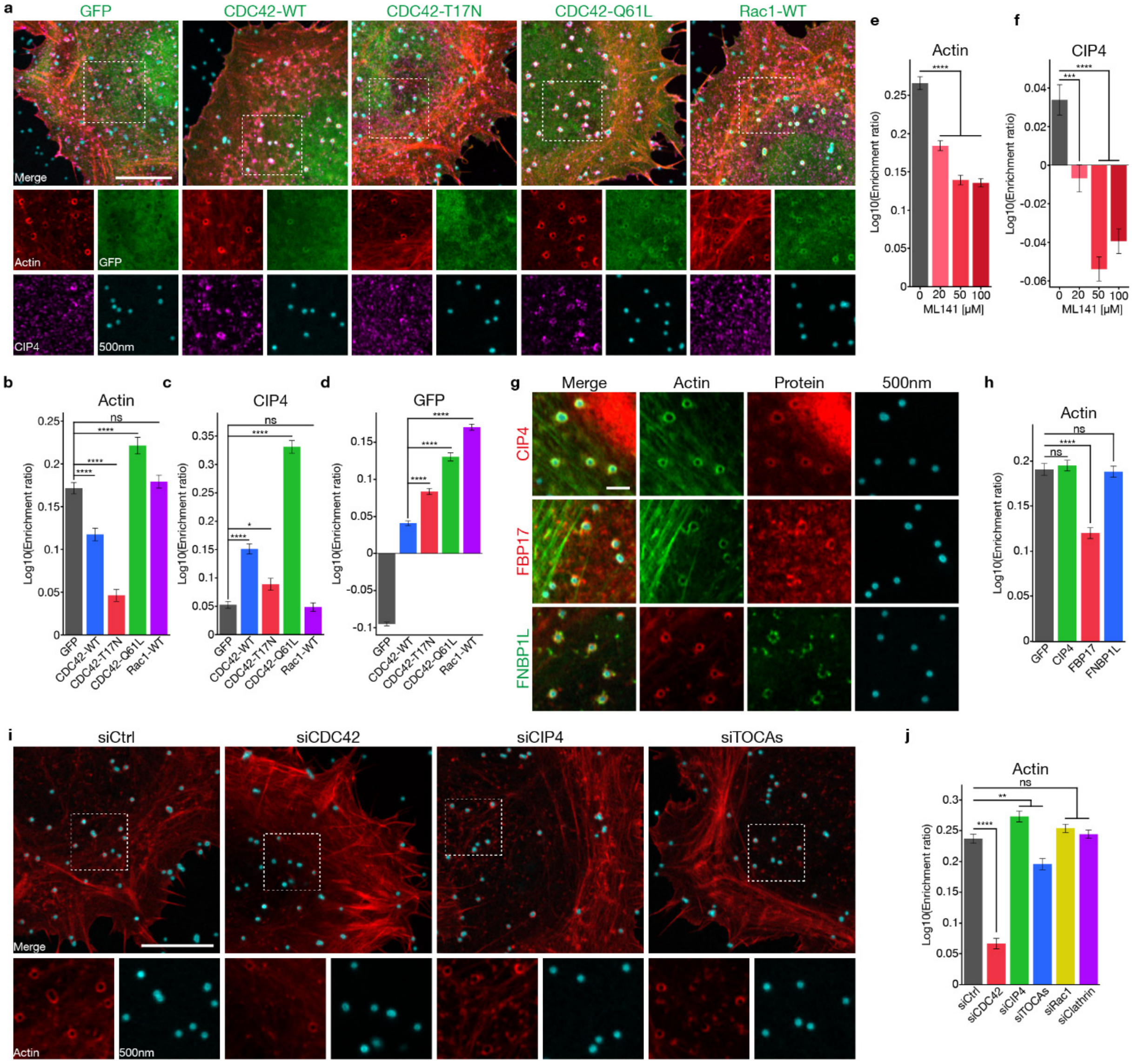


**Supplementary Figure 5: CDC42 and CIP4 control actin polymerisation around 500 nm plasma membrane deformations.** HeLa cells grown on 500 nm nanostructures and transfected with fluorescent constructs (**a-d,g-h**), treated with a specific CDC42 inhibitor (ML141) (**e-f**) or with siRNAs (**i-j**), as indicated. Data shown are quantifications of actin (**b,e,h,j**), endogenous CIP4 (**c,f**) or GFP (**d**) fluorescence around 500 nm deformations and the corresponding representative Airyscan images (**a,g,i**), as indicated. **a-d**, Effect of transient expression of GFP, GFP-CDC42-WT, GFP-CDC42-T17N, GFP-CDC42-Q61L or GFP-Rac1-WT, on the enrichment of actin, CIP4 and GFP around deformations. Number of deformations: GFP, n = 1379; CDC42-WT, n = 1214; CDC42-T17N, n = 602; CDC42-Q61L, n = 801; Rac1-WT, n = 997. Three independent experiments. **e-f**, Effect of dose-dependent inhibition of CDC42 by ML141 (0 μM, 20 μM, 50 μM or 100 μM) on the enrichment of actin and CIP4 around deformations. Number of deformations: 0 μM, n = 867; 20 μM, n = 1212; 50 μM, n = 1168; 100 μM, n = 1384. Three independent experiments. **g-h**, Effect of transient expression of GFP or individual TOCA proteins (mCherry-CIP4, mCherry-FBP17 or GFP-FNBP1L) on the enrichment of actin around deformations. Number of deformations: GFP, n = 1636; CIP4, n = 1995; FBP17, n = 1410; FNBP1L, n = 1556. Four independent experiments. **i-j**, Effect of the depletion of CDC42, CIP4, TOCAs (CIP4, FBP17 and FNBP1L), Rac1 or Clathrin Heavy Chain with siRNAs on the enrichment of actin around deformations. Number of deformations: siCtrl, n = 1238; siCDC42, n = 470; siCIP4, n = 789; siTOCA, n = 834; siRac1, n = 1562; siClathrin, n = 1417. Three independent experiments. Data are mean ± s.e.m.; ns, not significant; **** $P < 0.0001$; *** $P < 0.001$; ** $P < 0.01$; * $P < 0.05$ (one-way ANOVA with Dunnett's multiple comparison test). Regions marked by dashed squares, expanded below with individual channels displayed (**a,i**, bottom). White arrowheads, co-localisation (**a,g,i**). Scale bars, 10 μm (**a,i**) and 2 μm (**g**).

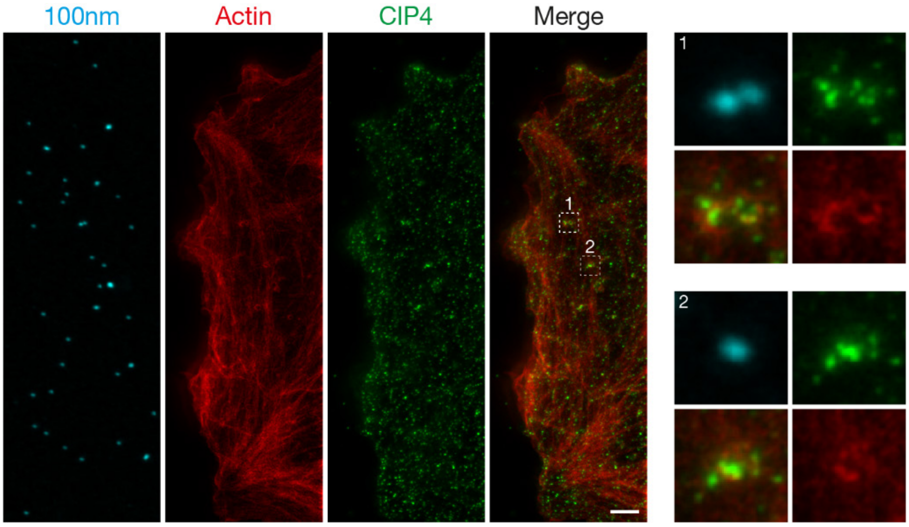


**Supplementary Figure 6: CIP4 and actin are specifically recruited to 100 nm deformations.** STED microscopy of cells grown on 100 nm nanostructures (cyan) and labeled for actin (phalloidin, red) and CIP4 (green). CIP4 and actin channels imaged in STED mode, nanostructures imaged in confocal mode. Regions #1 and #2 marked by dashed squares, expanded on the right with individual channels displayed. Scale bar, 2 µm.

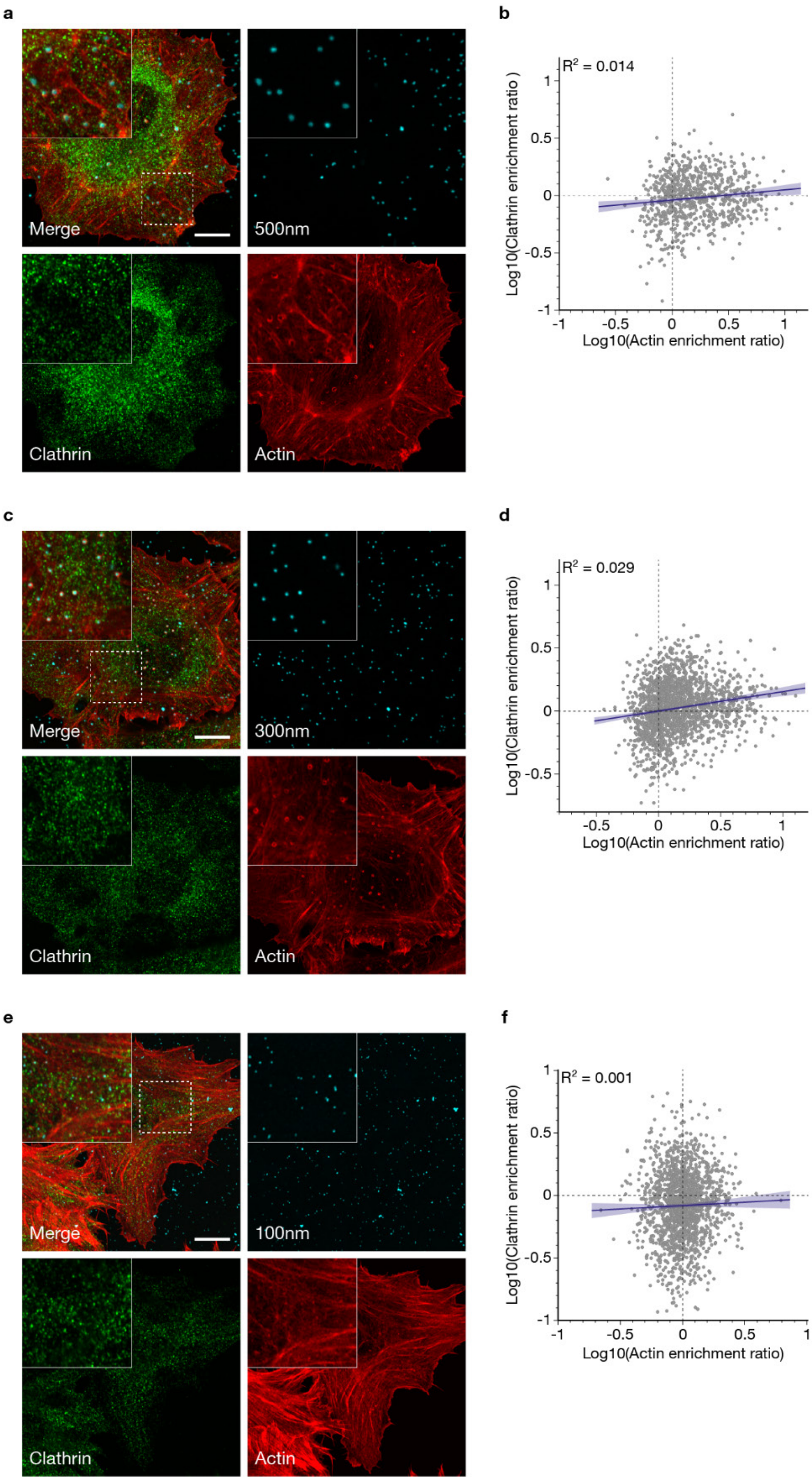


**Supplementary Figure 7: Curvature-induced actin polymerisation is independent of clathrin recruitment.** Actin and clathrin enrichment around 500 nm (**a-b**), 300 nm (**c-d**) and 100 nm (**e-f**) deformations. **a,c,e**, Representative Airyscan images. **b,d,f**, Correlations between the enrichment ratios of clathrin and actin around deformations. Number of deformations: 500 nm, n = 802; 300 nm, n = 1918; 100 nm, n = 1715. Linear regression with 95% confidence interval (blue). Three independent experiments. Insets, zoom on the dashed square regions. Scale bars, 10 µm.

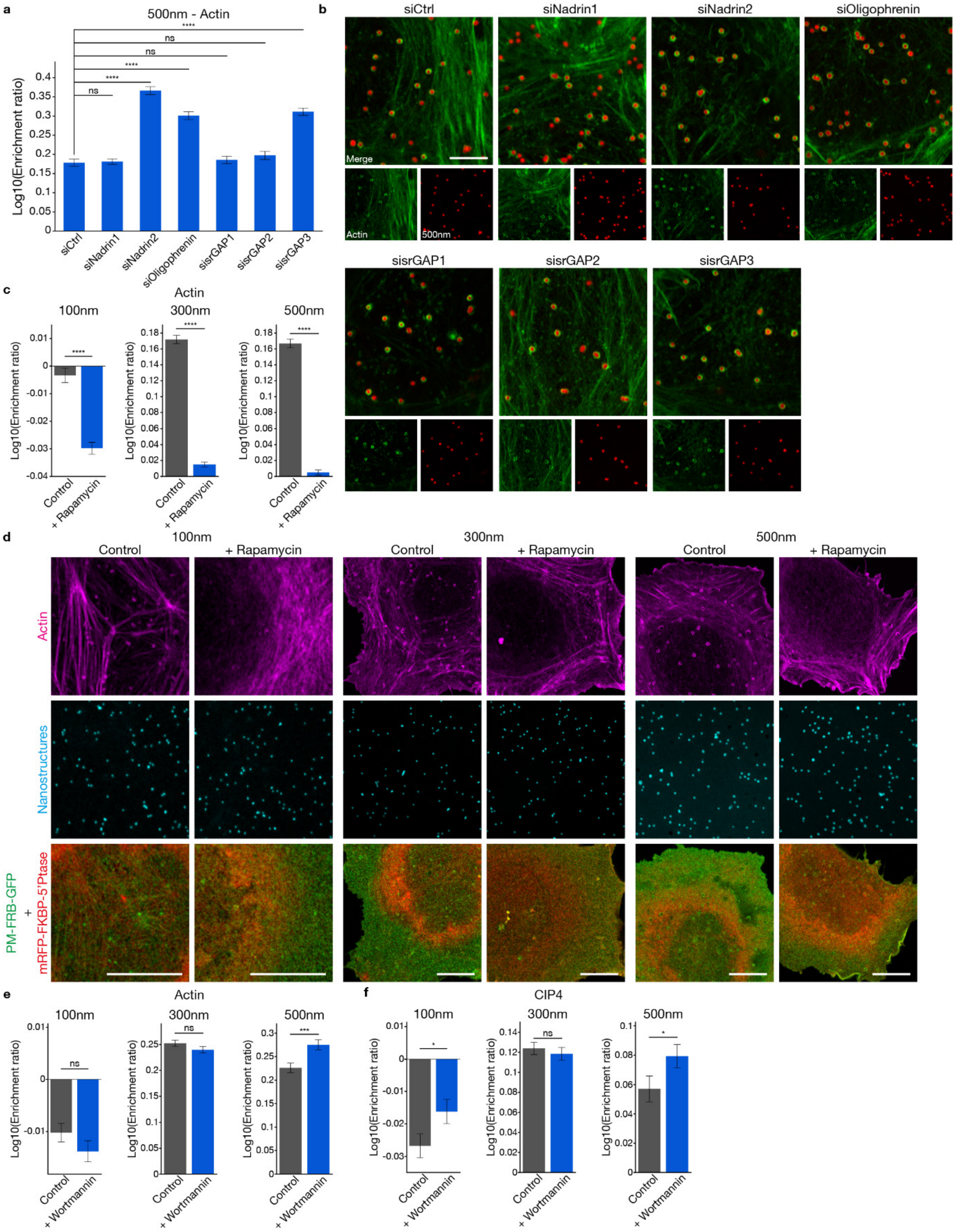


**Supplementary Figure 8: Rho GAP domain-containing BAR domain proteins and phosphoinositides regulate actin polymerisation around plasma membrane deformations. a-b**, Actin enrichment around 500 nm plasma membrane deformations upon Rho GAP domain-containing BAR domain proteins depletion with siRNAs. Quantifications of enrichment ratios from Airyscan images (**a**). Representative images (**b**). Number of deformations: siCtrl, n = 748; siNadrin1, n = 1245; siNadrin2, n = 811; siOligophrenin, n = 677; sisrGAP1, n = 688; sisrGAP2, n = 425; sisrGAP3, n = 940. Two independent experiments. **c-d**, Effect of PI(4,5)$P_2$ depletion on actin recruitment to deformations. Quantifications of enrichment ratios (**c**) and representative images (**d**) of actin around plasma membrane deformations before (Control) and after (+ Rapamycin) rapamycin (20 nM) addition inducing the recruitment of a 5'Ptase to plasma membrane. Number of deformations: 100 nm/Control, n = 3932; 100 nm/+ Rapamycin, n = 4015; 300 nm/Control, n = 1753; 300 nm/+ Rapamycin, n = 2208; 500 nm/Control, n = 1843; 500 nm/+ Rapamycin, n = 1835. Three independent experiments. **e-f**, Effect of PI3P species depletion on actin recruitment to deformations. Actin (**e**) and CIP4 (**f**) enrichment around plasma membrane deformations when PI3Ps are depleted from cells by inhibiting PI3 kinases with wortmannin (1 µM). Number of deformations: 100 nm/Control, n = 7210; 100 nm/+ Wortmannin, n = 6105; 300 nm/Control, n = 1745; 300 nm/+ Wortmannin, n = 1580; 500 nm/Control, n = 711; 500 nm/+ Wortmannin, n = 711. Three independent experiments. Data are mean ± s.e.m.; ns, not significant. **** $P < 0.0001$; *** $P < 0.001$; * $P < 0.05$ (**a**, one-way ANOVA with Dunnett's multiple comparison test; **c,e-f**, two-tailed unpaired t tests). Scale bars, 5 µm (**b**) and 10 µm (**d**).

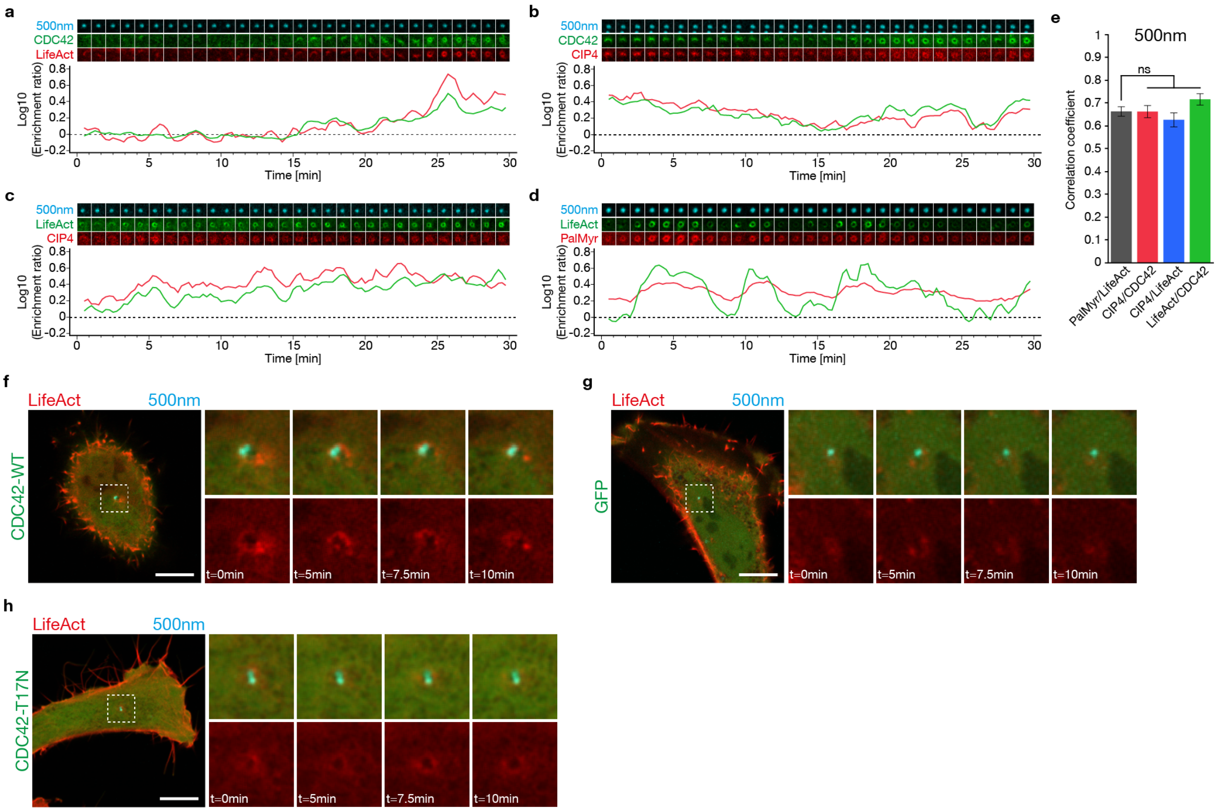


**Supplementary Figure 9: Local CIP4/CDC42-dependent actin polymerisation is dynamic and promoted by plasma membrane curvature. a-d**, Representative Airyscan time-lapses and the corresponding quantifications of enrichment ratios of the following pairs around 500 nm deformations over time: **a**, GFP-CDC42 and LifeAct-mCherry; **b**, GFP-CDC42 and mCherry-CIP4; **c**, LifeAct-GFP and mCherry-CIP4; **d**, LifeAct-GFP and PalMyr-mCherry (PalMyr). 30 min time-lapses with 15 sec intervals between frames. **e**, Correlation coefficients between the enrichment ratios over time curves for the pairs shown in **a-d**. Number of deformations: PalMyr/LifeAct, n = 99; CIP4/CDC42, n = 60; CIP4/LifeAct, n = 43; LifeAct/CDC42, n = 69. Three independent experiments. **f-h**, Monitoring of CDC42-dependent actin polymerisation upon deformation of the plasma membrane by FluidFM. **f-g**, Representative images of FluidFM/confocal experiments upon co-expression of: **f,** LifeAct-mCherry and GFP-CDC42-WT; **g,** LifeAct-mCherry and GFP; **h,** LifeAct-mCherry and GFP-CDC42-T17N. 10 min time-lapses with 9 sec intervals between frames. Two independent experiments. Regions marked by dashed squares surrounding the beads, expanded below at the indicated time points; t = 0 min, initial deformation of the cell membrane. Data are mean ± s.e.m.; ns, not significant (**e**, one-way ANOVA with Dunnett's multiple comparison test). Scale bar, 10 μm (**f-h**).

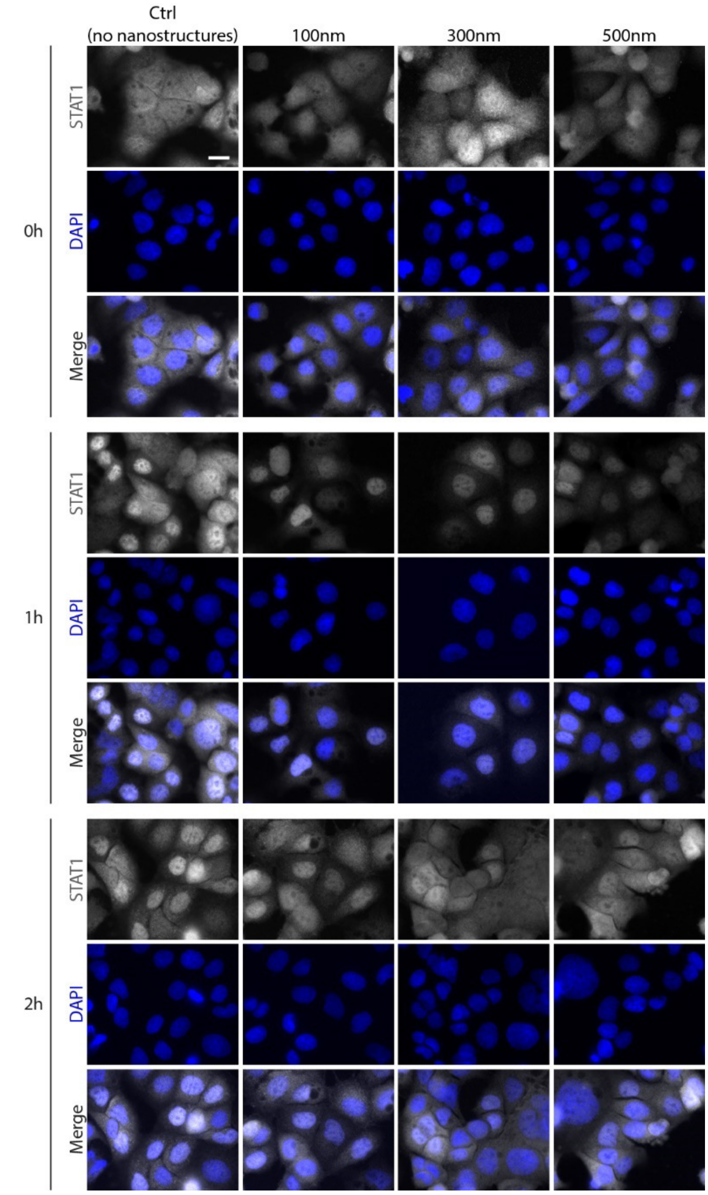


**Supplementary Figure 10: Representative images of STAT1 translocation experiments (related to Figure 5c-d).** Representative images of two independent experiments for 100 nm nanostructures and three independent experiments for other conditions. 0h, no IFNγ stimulation. Scale bar, 20 µm.

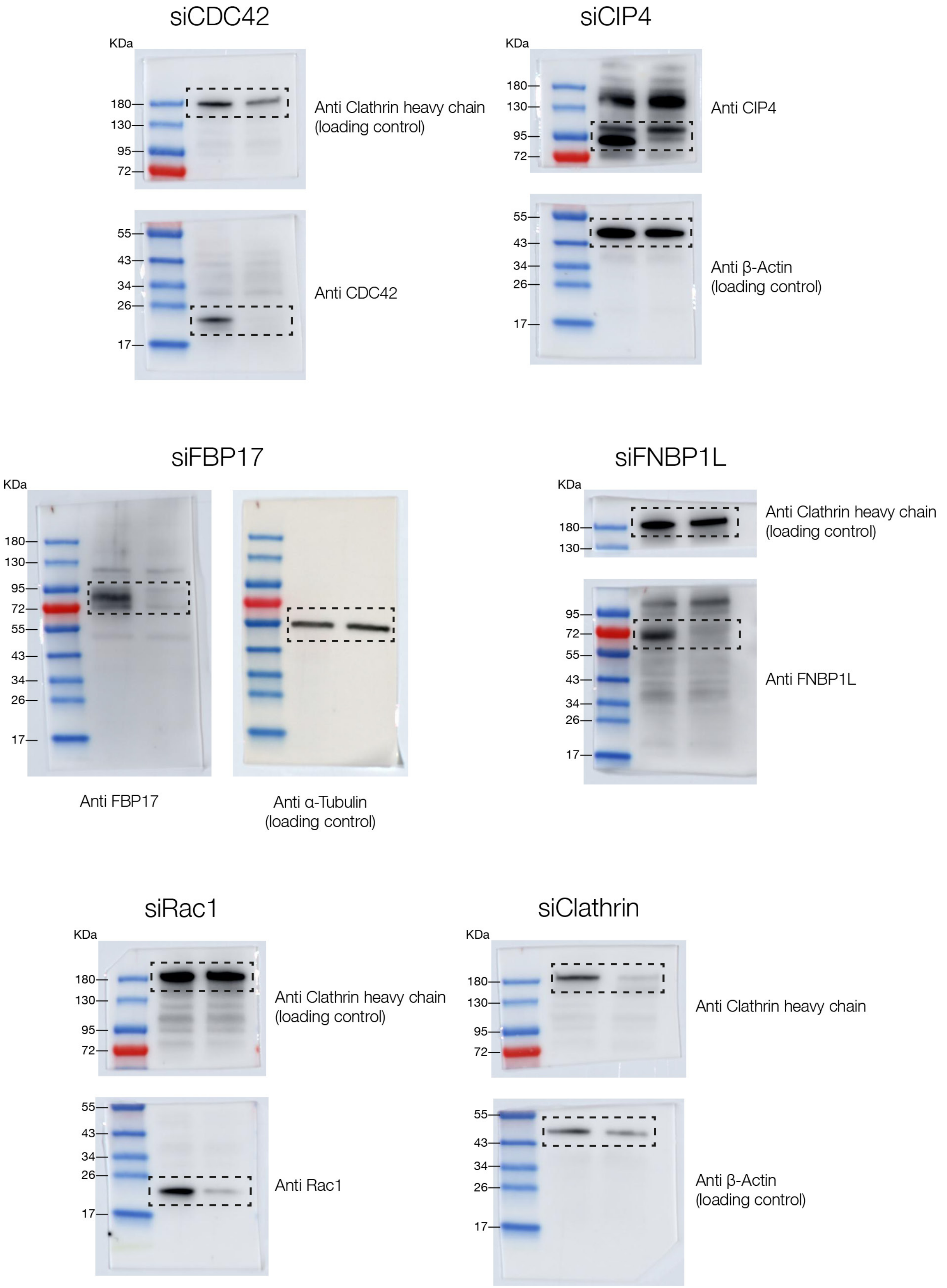


**Supplementary Figure 11: Unedited Western blot images of Figure 3, with molecular weights.** Dashed black rectangles indicate cropped areas.

### Movie legends

**Movie 1: Nanostructures are stable and are not displaced nor internalized by cells.** HeLa cells transiently expressing LifeAct-GFP (green) imaged for 2 hours after seeding on 500 nm nanostructures (red). 1 min intervals between frames. Single experiment.

**Movie 2: Actin is dynamic around membrane deformations.** HeLa cells transiently expressing LifeAct-GFP (green) seeded on 500 nm nanostructures (red). 30 min time-lapse with 15 sec intervals between frames. Representative of > 3 independent experiments.

**Movie 3: CDC42 and LifeAct follow similar dynamics on 100 nm membrane deformations, related to Figure 4a.** HeLa cells transiently expressing GFP-CDC42 (green) and LifeAct-mCherry (red) seeded on 100 nm nanostructures (cyan). 30 min time-lapse with 15 sec intervals between frames. Representative of 3 independent experiments.

**Movie 4: CDC42 and CIP4 follow similar dynamics on 100 nm membrane deformations, related to Figure 4b.** HeLa cells transiently expressing GFP-CDC42 (green) and mCherry-CIP4 (red) seeded on 100 nm nanostructures (cyan). 30 min time-lapse with 15 sec intervals between frames. Representative of 3 independent experiments.

**Movie 5: LifeAct and CIP4 follow similar dynamics on 100 nm membrane deformations, related to Figure 4c.** HeLa cells transiently expressing LifeAct-GFP (green) and mCherry-CIP4 (red) seeded on 100 nm nanostructures (cyan). 30 min time-lapse with 15 sec intervals between frames. Representative of 3 independent experiments.

**Movie 6: LifeAct and PalMyr have independent dynamics on 100 nm membrane deformations, related to Figure 4d.** HeLa cells transiently expressing LifeAct-GFP (green) and PalMyr-mCherry (red) seeded on 100 nm nanostructures (cyan). 30 min time-lapse with 15 sec intervals between frames. Representative of 3 independent experiments.

**Movie 7: CDC42 and LifeAct follow similar dynamics on 500 nm membrane deformations, related to Supplementary Figure 9a.** HeLa cells transiently expressing GFP-CDC42 (green) and LifeAct-mCherry (red) seeded on 500 nm nanostructures (cyan). 30 min time-lapse with 15 sec intervals between frames. Representative of 3 independent experiments.

**Movie 8: CDC42 and CIP4 follow similar dynamics on 500 nm membrane deformations, related to Supplementary Figure 9b.** HeLa cells transiently expressing GFP-CDC42 (green) and mCherry-CIP4 (red) seeded on 500 nm nanostructures (cyan). 30 min time-lapse with 15 sec intervals between frames. Representative of 3 independent experiments.

**Movie 9: LifeAct and CIP4 follow similar dynamics on 500 nm membrane deformations, related to Supplementary Figure 9c.** HeLa cells transiently expressing LifeAct-GFP (green) and mCherry-CIP4 (red) seeded on 500 nm nanostructures (cyan). 30 min time-lapse with 15 sec intervals between frames. Representative of 3 independent experiments.

**Movie 10: LifeAct and PalMyr follow similar dynamics on 500 nm membrane deformations, related to Supplementary Figure 9d.** HeLa cells transiently expressing LifeAct-GFP (green) and PalMyr-mCherry (red) seeded on 500 nm nanostructures (cyan). 30 min time-lapse with 15 sec intervals between frames. Representative of 3 independent experiments.

**Movie 11: FluidFM confocal experiment with wild-type CDC42 expressing cells, related to Figure 4f and Supplementary Figure 9f.** HeLa cells transiently expressing GFP-CDC42-WT (green) and LifeAct-mCherry (red) were deformed by a 500 nm particle (cyan) immobilized at the tip of a FluidFM probe. 10 min time-lapse with 9 sec intervals between frames. Representative of 2 independent experiments.

**Movie 12: FluidFM confocal experiment with GFP expressing cells, related to Figure 4f and Supplementary Figure 9g.** HeLa cells transiently expressing GFP (green) and LifeAct-mCherry (red) were deformed by a 500 nm particle (cyan) immobilized at the tip of a FluidFM probe. 10 min time-lapse with 9 sec intervals between frames. Representative of 2 independent experiments.

**Movie 13: FluidFM confocal experiment with CDC42-T17N expressing cells, related to Figure 4f and Supplementary Figure 9h.** HeLa cells transiently expressing GFP-CDC42-T17N (green) and LifeAct-mCherry (red) were deformed by a 500 nm particle (cyan) immobilized at the tip of a FluidFM probe. 10 min time-lapse with 9 sec intervals between frames. Representative of 2 independent experiments.

**Movie 14: FluidFM confocal experiment with CDC42-Q61L expressing cells, related to Figure 4f-g.** HeLa cells transiently expressing GFP-CDC42-Q61L (green) and LifeAct-mCherry (red) were deformed by a 500 nm particle (cyan) immobilized at the tip of a FluidFM probe. 10 min time-lapse with 9 sec intervals between frames. Representative of 2 independent experiments.
